# A large chromosomal inversion shapes gene expression in seaweed flies (*Coelopa frigida*)

**DOI:** 10.1101/2021.06.03.446913

**Authors:** Emma L. Berdan, Claire Mérot, Henrik Pavia, Kerstin Johannesson, Maren Wellenreuther, Roger K. Butlin

**Author notes:** These authors contributed equally (Kerstin Johannesson, Maren Wellenreuther, Roger K. Butlin).

## Abstract

Inversions often underlie complex adaptive traits, but the genic targets inside them are largely unknown. Gene expression profiling provides a powerful way to link inversions with their phenotypic consequences. We examined the effects of the *Cf-Inv(1)* inversion in the seaweed fly *Coelopa frigida* on gene expression variation across sexes and life stages. Our analyses revealed that *Cf-Inv(1)* shapes global expression patterns but the extent of this effect is variable with much stronger effects in adults than larvae. Furthermore, within adults, both common as well as sex specific patterns were found. The vast majority of these differentially expressed genes mapped to *Cf-Inv(1)*. However, genes that were differentially expressed in a single context (i.e. in males, females or larvae) were more likely to be located outside of *Cf-Inv(1).* By combining our findings with genomic scans for environmentally associated SNPs, we were able to pinpoint candidate variants in the inversion that may underlie mechanistic pathways that determine phenotypes. Together the results in this study, combined with previous findings, support the notion that the polymorphic *Cf-Inv(1)* inversion in this species is a major factor shaping both coding and regulatory variation resulting in highly complex adaptive effects.

## INTRODUCTION

Chromosomal inversions, pieces of the chromosome that have been flipped 180°, are structural variants that may encompass hundreds of genes but segregate together as a single unit due to suppressed recombination. Recombination between arrangements (i.e., orientations) is reduced in heterokaryotypes but proceeds freely in both homokaryotypes. This reduced recombination can shield adaptive allelic combinations from gene flow, facilitating evolutionary processes such as local adaptation [1–3], sex chromosome evolution [4–6] and speciation [1, 3, 7–10]. Inversions underlie major phenotypic polymorphisms in a wide variety of taxa, such as male reproductive morphs in the ruff, *Philomachus pugnax* [11, 12] and Müllerian mimicry wing patterns in the butterfly *Heliconius numata* [13]. However, the reduced recombination that allows inversions to have these profound effects also clouds signatures of selection on individual loci due to extreme linkage disequilibrium. This encumbers detection of the mechanistic pathways that generate phenotypic effects as well as identification of the underlying adaptive variants.

The linkage disequilibrium in inversions presents many challenges to identify adaptive variation. Since recombination between arrangements is rare, forward genetic approaches like QTL mapping or genome wide association studies are not feasible for variation that is fixed between arrangements [14]. Additionally, the reduced recombination and effective population size within the inverted region facilitates the accumulation of neutral and deleterious variation [15], increasing divergence between the arrangements and increasing the likelihood of detecting phenotype or environment associations with non-causative loci. Finally, larger inversions, such as the *lnv4m* inversion in *Zea mays*, may contain hundreds of genes that affect a wide variety of phenotypes that vary in their selective pressures [16].

Transcriptomic analysis offers a way to address the links between individual loci and the phenotypic effects of an inversion by uncovering functionally important variation in a way that is not hindered by linkage disequilibrium in natural populations or recombination suppression in controlled crosses. This is because (1) the phenotypic effects of inversions might be underlain in part by changes in gene expression, and (2) overlap between differentially expressed genes (from transcriptomic studies) and outlier SNPs (from genomic studies, i.e. loci associated with adaptive traits or ecological factors) facilitates the identification of candidate genes [17–19].

There are three major (non-exclusive) ways that inversions may affect gene expression. First, inversions may modify the epigenetic environment near their breakpoints [20, 21]. Second, breakpoints may change the relative positions of genes and their transcription regulators, changing expression patterns [22, 23]. Third, the linked variation within an inversion can contain *cis* or *trans* acting regulatory elements that can evolve independently in the two arrangements due to suppressed recombination between them [16, 24–26]. As variants within inversions are highly linked, it is difficult to distinguish between *cis* regulation and *trans*-acting loci in linkage disequilibrium with their targets. Here we focus on whether the differentially-expressed loci are contained within the inverted region (hereafter referred to as *cis* regulated for karyotype) or if the differentially expressed loci are located in other areas of the genome (hereafter referred to as *trans* regulated for karyotype). Overall, these effects on gene expression can be fixed, vary across life stages or sexes, or show plastic responses to the environment.

In this study, we investigated the effect of a large inversion on expression variation and combined this analysis with previously published population genomic data to identify putatively adaptive loci. We use the seaweed fly, *Coelopa frigida*, which inhabits “wrackbeds” (accumulations of decomposing seaweed) on North Atlantic shorelines. This fly has an inversion polymorphism on chromosome I (*Cf-Inv(1)* spanning 60% of chromosome 1 and 10% of the genome, corresponding to about 25MB) [27]. *Cf-Inv(1)* has two highly diverged arrangements, termed α and β, resulting from 3 overlapping inversions[28]. The inversion influences multiple measurable traits in males such as mating success [28–30], development time [31–33], longevity [34] and adult size [31, 35]. Of these, size is the trait where the inversion has the strongest effect; αα males are approximately three-fold heavier than ββ males [36]. This is mirrored in development time with αα males taking significantly longer to reach adult eclosion than ββ males [31]. Conversely, female phenotype is mostly unaffected by karyotype although there are small effects on size [33, 34]. The sex difference in the effect of the inversion indicates a particular role for gene expression as males and females largely share the same genome. The inversion is polymorphic in all investigated natural populations to date and maintenance of the polymorphism is mostly through balancing selection caused by strong overdominance of the heterokaryotype [29, 35, 37–39]. The spatial distribution of the inversion frequencies is then modulated by the seaweed composition and abiotic conditions of the wrackbed, which influence the relative fitnesses of the homokaryotypes [29, 35, 36, 40].

We collected *C. frigida* from natural populations (Figure 1A) and examined how *Cf-Inv(1)* shaped gene expression across sexes and life stages. Specifically, our study had three major goals: 1. To examine the role of the inversion in shaping global expression patterns in adults and larvae and to determine if these effects are common or context specific, 2. To ascertain if these genes are *cis* or *trans* regulated with respect to *Cf-Inv(1),* and 3. To identify putative adaptive variation within the inversion and connect this with ecological niche differences between karyotypes.

**Figure 1.**
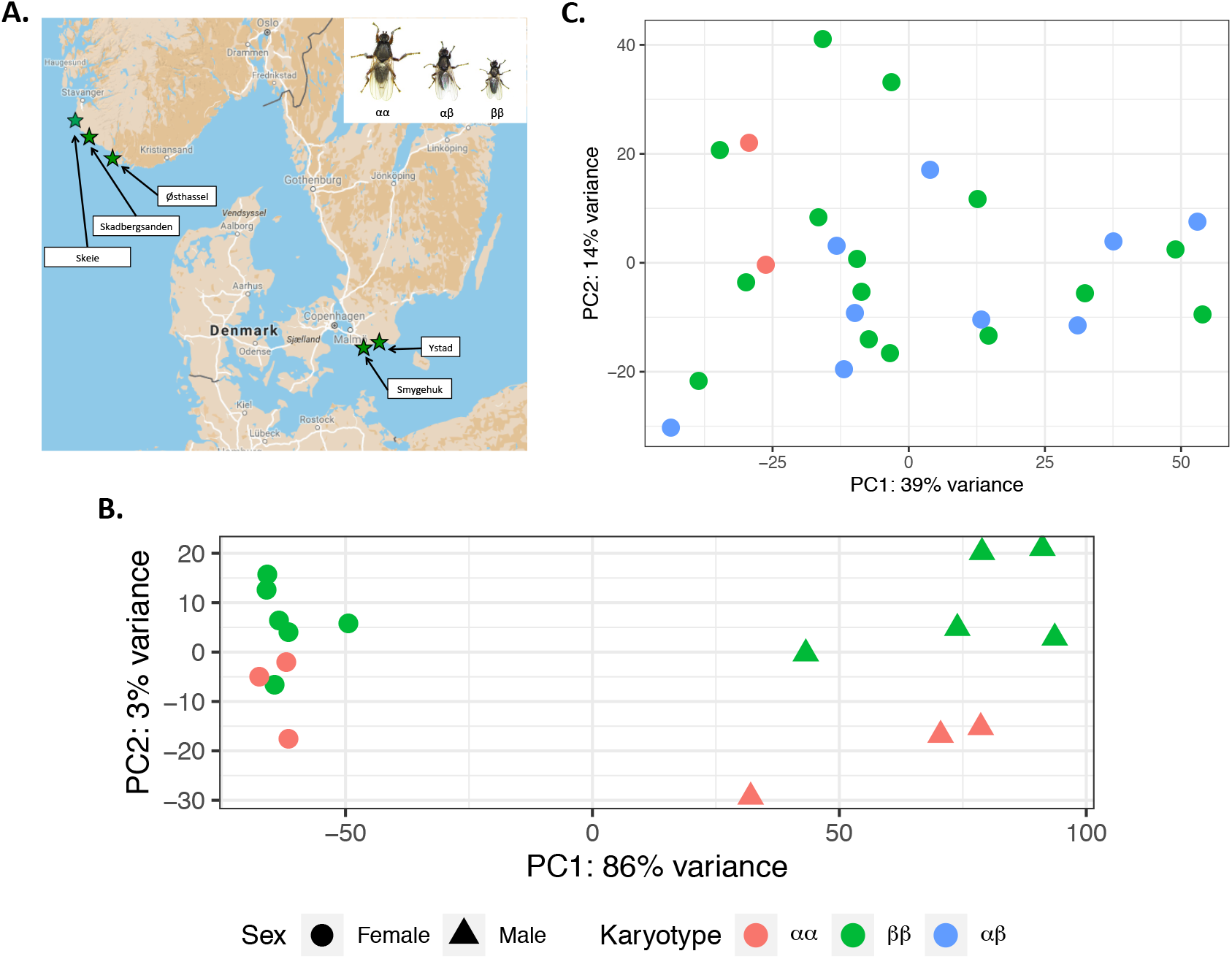
Variation in expression differs across life stages **A**. Map of Norway, Denmark, and Sweden showing the populations sampled. The inset shows size variation in males as a function of karyotype. **B**. Principal component analysis (PCA) of expression variation in adults. Points are colored by karyotype (αα- red, ββ-blue) and shaped according to sex (female-circle, male-triangle). **C**. PCA of expression variation in larvae, all samples are pools of 3 larvae of unknown sex colored by karyotype (αα- red, αβ-green, ββ-blue). Both Figure 1A and 1B are based on the top 500 transcripts with the highest variance.

## RESULTS AND DISCUSSION

### Sequencing and transcriptome assembly

To study gene expression variation associated with sex, life stage and karyotypes of the inversion, we sequenced RNA from 17 adult individuals and 28 larval pools. We used part of this data set to create the first reference transcriptome for *C. frigida*. Our final transcriptome assembly contained 35,999 transcripts with an N50 of 2,155bp, a mean length of 1,092bp, and a transrate score [41] of 0.4097. The transcriptome has good coverage, it has a BUSCO score of 86.6% (2,393 complete and single copy (85.5%), 31 complete and duplicated (1.1%), 190 fragmented (6.8%), and 185 missing (6.6%)), and 95% of the reads mapped back to the transcriptome [42]. Using the trinotate pipeline, we were able to partially annotate 14,579 transcripts (40%) from the transcriptome. This high-quality transcriptome will provide a useful resource for any future work on this and related species, provide a much needed functional map for better understanding the regulation of genes across life stages and sexes, and facilitate the identification of functional phenotypes that correspond to inversions.

### The effect of *Cf-Inv(1)* on gene expression is strong but variable

In adults, karyotype was the second strongest factor explaining expression variation. Decomposing adult expression variation into a PCA we found that the PC1, explaining 86% of the variance, separated males and females while PC2, explaining 3% of the variance, separated αα and ββ in both males and females (Figure 1B). This strong sex difference was mirrored in our differential expression analysis; a total of 3,526/26,239 transcripts were differentially expressed between the sexes with a strong bias towards increased expression in males (68% of differentially expressed genes upregulated in males, Supplemental Figure 1).

Sex modulated the effects of *Cf-Inv(1)* on global expression patterns. When combining the sexes, 304/26,239 transcripts were differentially expressed between αα and ββ (Supplemental Figure 2). A distance matrix analysis revealed that (1) average similarity between pairs of females was higher than between pairs of males and (2) males clustered by karyotype while females did not (Supplemental Figure 3). Due to these strong differences we chose to run separate analyses for the sexes instead of analyzing the interaction term from our main model. Comparing homokaryotypic sex groups separately (αα vs. ββ) revealed that more than double the number of differentially expressed genes were found in males compared to females (801 vs 340; Supplemental Figure 4, 5). There was substantial overlap between these groups with the highest proportion of unique differentially expressed genes found in males (Supplemental Figure 6). The phenotypic effects of *Cf-Inv(1)* are also strongly sex-specific. This is likely due to sexual selection which, in *C. frigida*, has partly evolved in response to strong sexual conflict over reproduction, specifically mating rate [43, 44]. This sexual conflict over mating rates has selected for sexual dimorphism in some of the external phenotypic traits used for mating, notably size. Larger males are more successful in obtaining copulations and resisting the rejection responses that females use to prevent male mountings. The *Cf-Inv(1)* inversion has a large impact on the morphology of males [31, 32]. It was thus no surprise that males showed a larger gene expression difference between karyotypes compared to females.

Surprisingly, *Cf-Inv(1)* was not a primary factor explaining variance in larval gene expression. A PCA in larvae found that the first two PCs (explaining 52% of the variance) did not separate samples based on karyotype (Figure 1C), instead a separation by population was observed (Supplemental Figure 7). We ran an additional PCA on the larval data using only the Skeie population (the only population with all 3 karyotypes), to remove population variation. The first two PCs (explaining 67% of the variance) together separated the karyotypes, albeit weakly (Supplemental Figure 8).

To formally test the role of karyotype in partitioning variation we ran a PERMANOVA on Manhattan distances for each subgroup (i.e. males, females, and larvae; Supplemental Table 2)[45]. As different tests had different sample sizes, we concentrated on R^2^ values (sum of squares of a factor/total sum of squares). Males and females had the highest R^2^ values (0.2464 and 0.153, respectively) followed by all adults and larvae (0.084 and 0.073, respectively). These results match our qualitative observations that karyotype explains the largest proportion of variance in adult males followed by adult females and then larvae. However, the comparison of our combined adult model with the sex specific models shows that separating sex is critical for quantifying the effect of karyotype. Thus, the superficial appearance of inversion having less influence on larval gene expression may be because larval sex was not determined.

Further dissecting differential expression in our full larval data set corroborated our qualitative observations. Since we had three genotypes in larvae (αα, αβ, and ββ) we ran three different contrast statements (αα vs. ββ, αβ vs. ββ, and αα vs. αβ). When comparing expression in ββ vs. αβ we found that 23/15,859 transcripts were differentially expressed and most of these (74%) were upregulated in αβ (Supplemental Figure 9). Comparing expression in αα vs. ββ, we found 29/15,859 transcripts to be differentially expressed and most of these (83%) were upregulated in ββ (Supplemental Figure 10). Comparing expression in αα vs. αβ, we found 6/15,859 transcripts to be differentially expressed and most of these (83%) were upregulated in αβ. There was some overlap between these three contrasts (Supplemental Figure 11). Overall, a greater portion of transcripts were significantly differentially expressed between αα vs. ββ in adults (1.16%-3.05%) compared to larvae (0.2%). In addition to pooling sexes in larvae there are several other features of our experimental design that could have contributed to the reduced effect in larvae. First, our crossing-design generated only two αα larval pools compared to ten αβ larval pools and sixteen ββ larval pools. Thus, our contrasts that included αα had lower power. We also generated more variation in our larval samples compared to our adults as we crossed both within and between populations while adults were all single population origin. It is possible that this variation made detection of differentially expressed genes more difficult. However our results still clearly suggest that the effect of *Cf-Inv(1)* on gene expression is strongly conditional on life stage and sex.

### Allele specific expression within *Cf-Inv(1)*

Beyond quantitative differences of expression, genes within *Cf-Inv(1)* were also characterized by allele-specific expression (ASE) in heterokaryotes. Concentrating on loci that were fixed between arrangements, we retained 315/619,424 SNPs found across 113 transcripts all located within *Cf-Inv(1).* Using the ASEP package [46] with our 9 αβ larval pools, a total of 30/113 transcripts had significant ASE (Supplemental Figure 12). We compared this with our complete differential expression results and found that only a single transcript overlapped between the two. For each of these transcripts we averaged read depth across all SNPs per transcript, per individual. We classified them as ‘α biased expression’ if > 50% of the larval pools had >=55% α-allele reads and as ‘β biased expression’ if > 50% of the larval pools had >=55% β -allele reads. If neither of these conditions was met, i.e. the direction was inconsistent; we simply labeled them as ‘allele biased expression’. We found 5 transcripts that showed ‘α biased expression’, 12 transcripts that showed ‘β biased expression’, and 13 transcripts that showed ‘allele biased expression’ (Supplemental Figure 13, transcripts with data for => 5 individuals is shown in Figure 2). There were no significant GO terms for any of these groupings. Two interesting patterns emerge from these data. First, allele biased expression, when present, seems to be relatively consistent across populations. Our αβ larvae resulted from crosses within and between populations yet we found consistent ASE patterns in 56% of our ASE transcripts. Second, differentially expressed genes showed no propensity towards ASE as only 1/30 ASE genes showed significant differential expression and most showed close to zero differential expression (e.g., from the combined adult αα vs. ββ comparison the mean absolute log2Fold change was 0.75). This indicates that ASE may be evolving somewhat independently from differential expression. Overall, these results demonstrate that there is allele biased expression within inversions but the extent of this phenomenon and the resulting phenotypic implications remain unknown.

**Figure 2.**
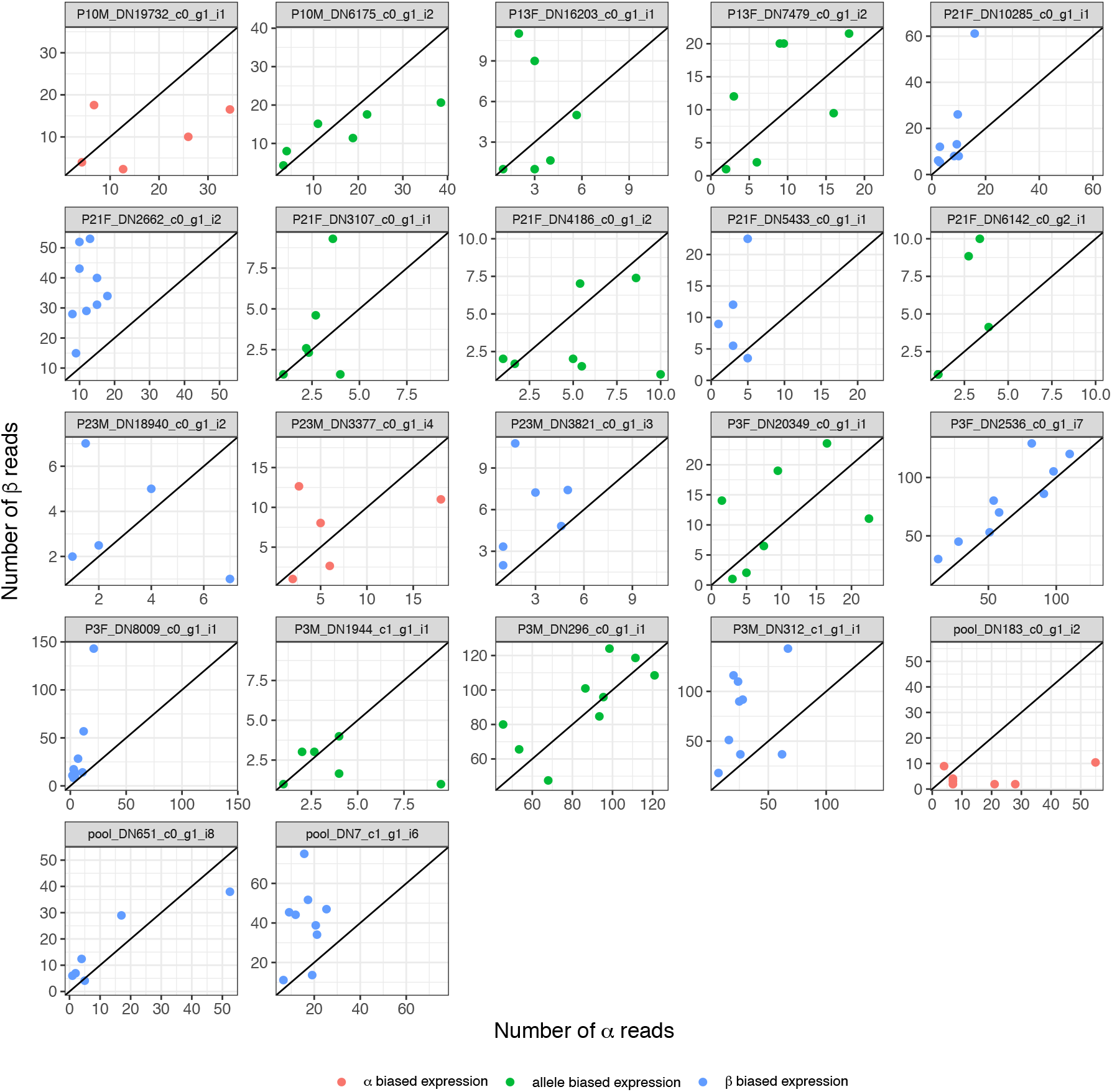
Patterns of allele specific expression (ASE). Each plot is for a single transcript where each dot represents a single αβ individual averaged over all SNPs in that transcript. A 1:1 line is provided for context. Colors indicate the expression pattern: α biased expression – red, β biased expression – blue, allele-biased expression – green. Note that only transcripts with data for 5 or more individuals are shown here. The full data set is shown in Supplemental Figure 12.

### Genes with constant karyotype effects are overwhelmingly *cis*-regulated while genes with conditional effects are more likely to be *trans*-regulated

Most of the differentially expressed genes mapped within *Cf-Inv(1)* (Figure 3). For adults, 12.8% of transcripts tested for differential expression were found within *Cf-Inv(1)* (Table 1) which is approximately what might be expected, as *Cf-Inv(1)* comprises 10.5% of the genome [27]. However, 80.6% of the transcripts that were differentially expressed between αα and ββ (with the sexes combined) were found within *Cf-Inv(1)* (odds ratio = 28.3). Looking at this in a different way, 7.2% of the transcripts within the inversion were differentially expressed between karyotypes compared to 0.3% of genes in the collinear region. When decomposing the sexes, the *cis*-effect was much stronger in females than males as 78% of differentially expressed genes in females (odds ratio = 24.2) were found within *Cf-Inv(1)* compared to 44.5% in males (odds ratio = 5.5; Figure 3A,B). For larvae we combined the ββ vs. αβ, αα vs. αβ and αα vs. ββ contrasts as so few differentially expressed transcripts were found (a combined total of 55 transcripts). Of these, 52.8% were found within *Cf-Inv(1)* (odds ratio = 7.6). This effect is visible when comparing density plots for log2fold changes from αα vs. ββ comparisons from the entire genome to within *Cf-Inv(1)* (Figure 3B,D,F). Here we see two trends. First the whole genome density plots for both males (Figure 3B) and larvae (Figure 3F) are much flatter and left shifted than the density plot for females (Figure 3B). Second, for all three groups the density plots for genes within *Cf-Inv(1)* are wider and more left-shifted. All of these differences were significant with two sample Kolmogorov-Smirnov tests but the effect was weaker when comparing the whole genome vs. within *Cf-Inv(1)* in larvae (Supplemental Table 3). Compared to karyotype the effect of sex showed no pattern of localization. Instead, transcripts differentially expressed between males and females in adults closely matched the null distribution of tested transcripts (Table 1).

**Figure 3.**
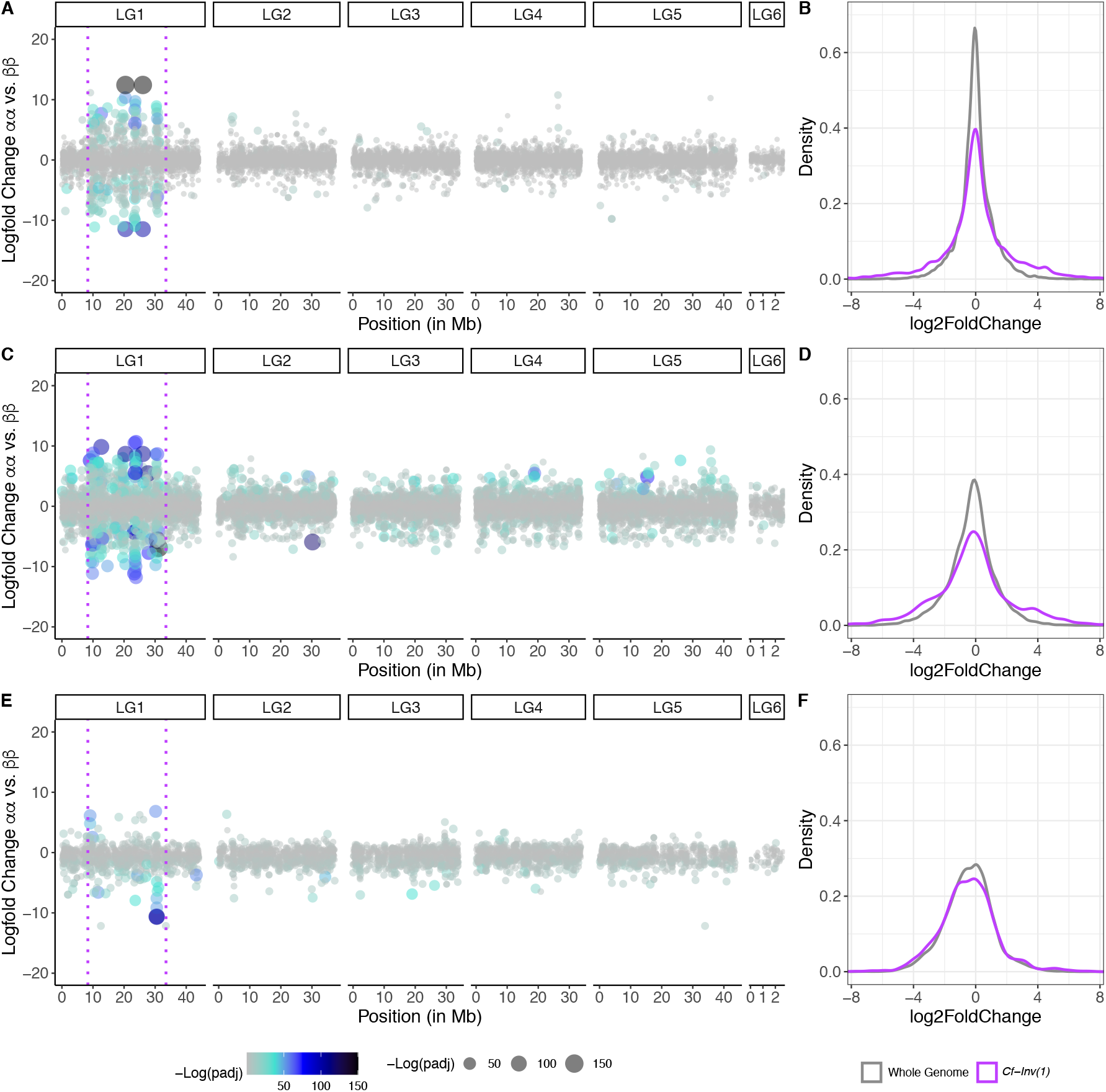
Differential expression is mostly *cis*-regulated for karyotype. Differentially expressed transcripts along the genome in (A) females, (C) males, and (E) larvae. Y-axes denote logfold change between αα and ββ and x-axes denote position in megabases. The dotted magenta lines denote the location of *Cf-Inv(1)*. Note that position in LG6 is not to scale with the other linkage groups for presentation. Each dot is a single transcript and both color and size denote the −log (p-value) after false discovery rate correction. Next to each graph are density plots of log2fold changes for αα vs. ββ comparisons for all loci in the genome (colored grey) and just loci in within *Cf-Inv(1)* (colored magenta) for each group: females (B), males (D) and larvae (F). Negative values indicate higher expression in ββ.

**Table 1.** Location of differentially expressed transcripts. Proportion of differentially expressed or tested transcripts are shown as percentages located within different linkage groups or inversions. The ‘Other Scaffolds’ category sums across 340 scaffolds that could not be incorporated into existing linkage groups [For details see 27]. The total number of transcripts represented by each group is: 25,320 (tested transcripts), 293 (DE between αα and ββ), and 3,411 (DE between males and females).

| Location | Tested Transcripts | Differentially Expressed between $\alpha\alpha$ and $\beta\beta$ | Differentially expressed between males and females |
| --- | --- | --- | --- |
| LG1 | 10.6% | 3.1% | 12.0% |
| *Cf-Inv(1)* | 12.8% | 80.5% | 11.6% |
| LG2 | 18.6% | 2.7% | 18.8% |
| LG3 | 16.6% | 3.8% | 17.4% |
| LG4 | 18.0% | 3.8% | 19.0% |
| LG5 | 17.5% | 3.4% | 17.6% |
| LG6 | 1.6% | 0.0% | 0.7% |
| Other Scaffolds | 4.3% | 2.7% | 2.9% |

The fact that most of the differentially expressed genes were *cis*-regulated for karyotype but not for sex effects is consistent with the idea that gene expression presents a major substrate for evolutionary change. For example, local adaptation via changes in gene expression has recently been demonstrated to be more common than by amino acid substitutions in humans [47]. Other recent studies of expression variation between karyotypes have also found strong *cis*-effects [22, 25, 26]. Allele biased expression is expected under *cis* regulation so these results are concordant with our ASE analysis [48]. Interestingly, the group where the strongest phenotypic differences are present (males) showed more *trans* effects of *Cf-Inv(1)*. Furthermore, differentially expressed transcripts that were shared between analyses were more likely to be located within *Cf-Inv(1).* Of transcripts significant in both the male and female comparisons, 92.3% map to *Cf-Inv(1)* compared with 59.8% of transcripts unique to the female analysis and 29% of transcripts unique to the male analysis. Overall, these results suggest that the ‘base’ effect of the inversion might be mostly *cis*-regulated while conditional effects may be more likely *trans*. *Cis-*regulatory elements are physically linked to the genes whose expression they control and thus tend to influence one or a few gene targets, often in specific tissues or at specific times, whereas more distant *trans* factors can control the expression of many genes. Thus, *trans* control of conditional effects in inversions may evolve more easily due to cascading effects. This is in line with evidence suggesting *trans* regulation may also be important for environment-dependent changes in gene expression [49, 50]. Our results highlight the importance of comparing the effects of inversions on gene expression in multiple contexts (i.e. sexes, life stages).

### Processes affected by *Cf-Inv(1)* include metabolism and development

To be able to connect changes in expression with the phenotypic effects of *Cf-Inv(1)* we first tested for enrichment of gene ontology (GO) categories in differentially expressed genes between karyotypes and sexes (Table 2). We found 16 significantly enriched GO terms across all of our tests but removed one GO term as it was supported by a single transcript. The 15 remaining terms can be found in Table 2. The three terms associated with karyotype related to development (adult chitin-based cuticle development) and metabolism/energy storage (digestion, positive regulation of triglyceride lipase activity). Unsurprisingly, the majority of the terms associated with sex differences were related to the production of gametes (e.g., sperm axoneme assembly, germ-line stem cell division).

**Table 2.** Significantly enriched Gene Ontology terms. Listed are: the GO ID, the term, the number of transcripts annotated with that term in the testing set, the number of these transcripts that were differentially expressed, the expected number of transcripts, the p-value from the elim model with Fisher Exact Test, the adjusted P-value, the analysis where the term was significant, and other analyses where the same term was significant. If a term was significant in multiple analyses we show the data from the most significant test and list that one in the analysis column.

| GO ID | Term | Annotated | Significant | Expected | elimF | Adjusted P-value | Analysis | Additional analyses where significant |
| --- | --- | --- | --- | --- | --- | --- | --- | --- |
| GO:0003341 | cilium movement | 59 | 35 | 10.14 | 1.00E-07 | 0.0003617 | Sex |  |
| GO:0006030 | chitin metabolic process | 96 | 32 | 16.5 | 8.60E-05 | 0.084835091 | Sex |  |
| GO:0006270 | DNA replication initiation | 26 | 18 | 4.47 | 6.30E-07 | 0.00162765 | Sex |  |
| GO:0007288 | sperm axoneme assembly | 15 | 11 | 2.58 | 2.60E-06 | 0.0047021 | Sex |  |
| GO:0007305 | vitelline membrane formation involved in chorion-containing eggshell formation | 19 | 15 | 3.27 | 6.20E-09 | 3.36E-05 | Sex |  |
| GO:0007586 | digestion | 99 | 10 | 1.03 | 7.00E-08 | 0.00075957 | Adult $\alpha\alpha$ vs. $\beta\beta$ | Female $\alpha\alpha$ vs. $\beta\beta$, Larvae $\alpha\alpha$ vs. $\beta\beta$ |
| GO:0008365 | adult chitin-based cuticle development | 8 | 7 | 0.25 | 1.90E-10 | 2.06E-06 | Male $\alpha\alpha$ vs. $\beta\beta$ | Sex |
| GO:0030720 | oocyte localization involved in germarium-derived egg chamber formation | 11 | 8 | 1.89 | 7.60E-05 | 0.0824676 | Sex |  |
| GO:0034587 | piRNA metabolic process | 22 | 13 | 3.78 | 1.20E-05 | 0.018601714 | Sex |  |
| GO:0035082 | axoneme assembly | 67 | 40 | 11.51 | 6.10E-09 | 3.36E-05 | Sex |  |
| GO:0042078 | germ-line stem cell division | 29 | 14 | 4.98 | 0.00011 | 0.0994675 | Sex |  |
| GO:0060294 | cilium movement involved in cell motility | 12 | 9 | 2.06 | 1.70E-05 | 0.023058375 | Sex |  |
| GO:0061365 | positive regulation of triglyceride lipase activity | 5 | 4 | 0.15 | 4.20E-06 | 0.0227871 | Male $\alpha\alpha$ vs. $\beta\beta$ | |
| GO:1905349 | ciliary transition zone assembly | 6 | 6 | 1.03 | 2.60E-05 | 0.031347333 | Sex |  |

We also investigated the impact of *Cf-Inv(1)* at the level of pathways by testing for polygenic expression patterns using Signet library [51]. We identified a number of gene subnetworks within biological pathways that show differential expression between karyotypes and sexes. Twenty-six pathways were differentially expressed between αα and ββ (Table 3A). Of these, 10 were found in multiple tests. We found pathways related to cell cycle metabolism and control, such as nucleotide metabolism or amino acid metabolism as well as signalling (FoxO pathway) or genetic information processing (Fanconi anemia pathway). Twelve of the 26 pathways differing between karyotypes were also related to energetic metabolism, particularly in males, including fatty acid degradation, carbohydrate metabolism and metabolism of co-factors. Of particular interest, male analysis included two organismal pathways, one related to longevity regulation and another involved in phototransduction in flies. As in other insects, increased size in *C. frigida* is associated with increased longevity and thus αα males live considerably longer on average [34]. We found 16 pathways differentially expressed between males and females (Table 3B), 9 of which were also identified in our karyotype analyses.

**Table 3.**
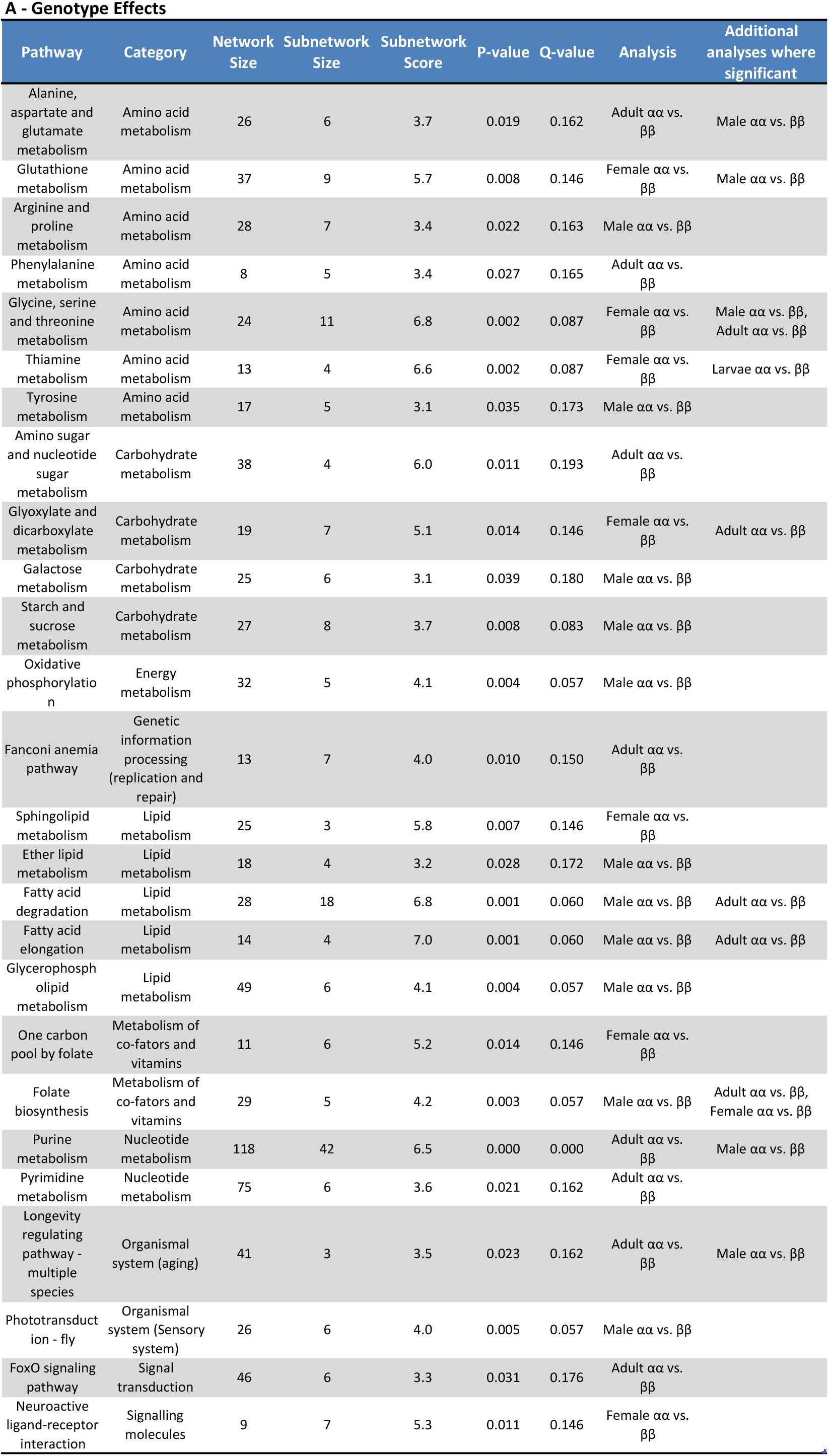
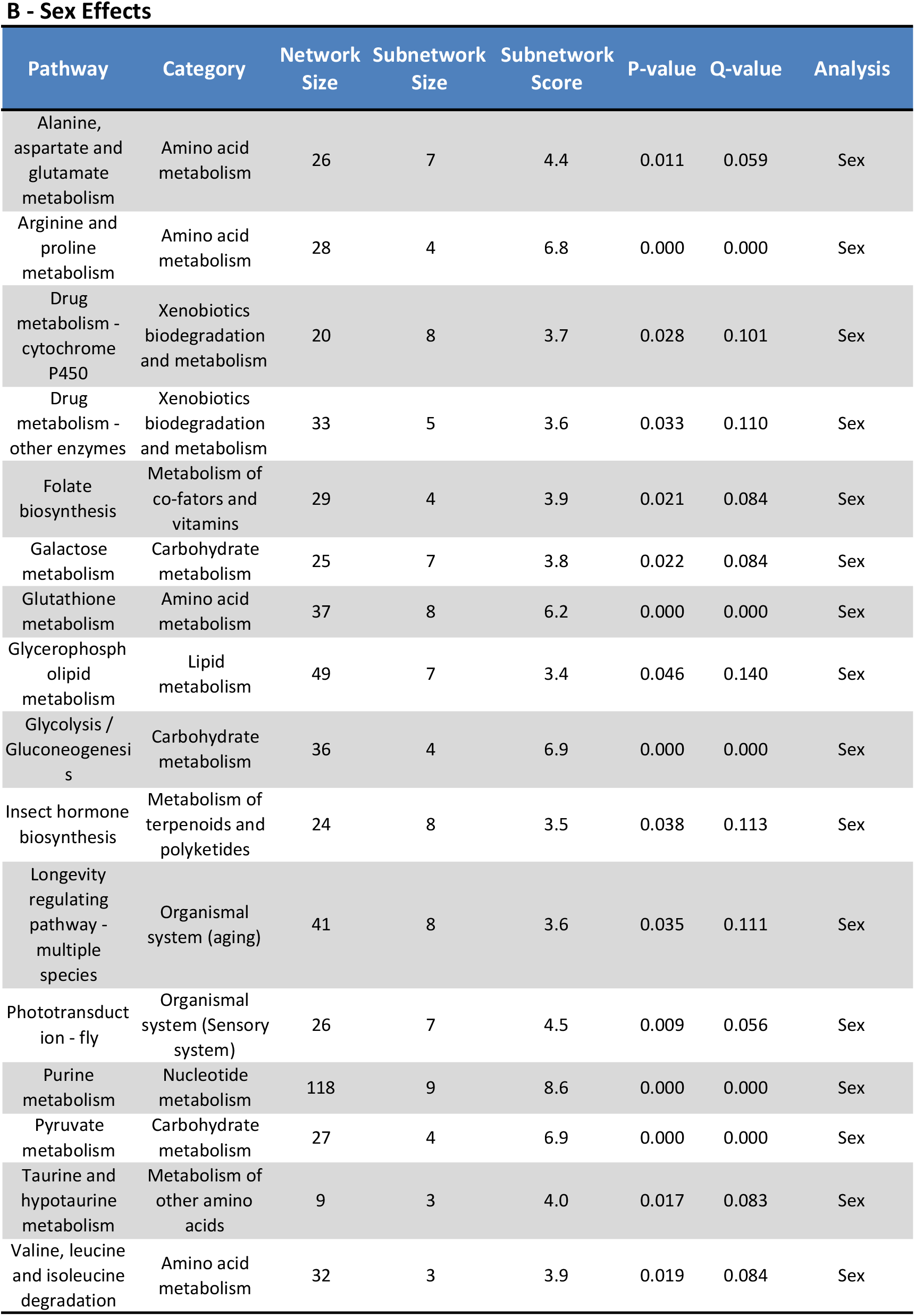
Functional pathways exhibiting subnetworks of genes interacting with each other and differentially expressed between karyotypes or sexes. For clarity, only karyotype effects are shown in (A) and Sex effects are shown in (B). Pathways are based on the KEGG database with genes identified in flybase. Significance of network score was assessed using the R library signet, by comparing to scores generated by random sampling. Network size is the number of genes connected in the pathways under consideration. Subnetworks are a subset of genes that are directly connected by edges and show high-scoring. Subnetwork size is the number of genes and subnetwork score is the normalized score inferred by the procedure based on the strength of the relationship between the factor compared (karyotype/sex) and expression at the genes involved in this subnetwork. For A, if a term was significant in multiple analyses we show the data from the most significant test and list that one in the analysis column. The additional tests are listed under ‘Additional analyses where significant’.

A - Genotype Effects
| Pathway | Category | Network Size | Subnetwork Size | Subnetwork Score | P-value | Q-value | Analysis | Additional analyses where significant |
| --- | --- | --- | --- | --- | --- | --- | --- | --- |
| Alanine, aspartate and glutamate metabolism | Amino acid metabolism | 26 | 6 | 3.7 | 0.019 | 0.162 | Adult $\alpha\alpha$ vs. $\beta\beta$ | Male $\alpha\alpha$ vs. $\beta\beta$ |
| Glutathione metabolism | Amino acid metabolism | 37 | 9 | 5.7 | 0.008 | 0.146 | Female $\alpha\alpha$ vs. $\beta\beta$ | Male $\alpha\alpha$ vs. $\beta\beta$ |
| Arginine and proline metabolism | Amino acid metabolism | 28 | 7 | 3.4 | 0.022 | 0.163 | Male $\alpha\alpha$ vs. $\beta\beta$ | |
| Phenylalanine metabolism | Amino acid metabolism | 8 | 5 | 3.4 | 0.027 | 0.165 | Adult $\alpha\alpha$ vs. $\beta\beta$ | |
| Glycine, serine and threonine metabolism | Amino acid metabolism | 24 | 11 | 6.8 | 0.002 | 0.087 | Female $\alpha\alpha$ vs. $\beta\beta$ | Male $\alpha\alpha$ vs. $\beta\beta$, Adult $\alpha\alpha$ vs. $\beta\beta$ |
| Thiamine metabolism | Amino acid metabolism | 13 | 4 | 6.6 | 0.002 | 0.087 | Female $\alpha\alpha$ vs. $\beta\beta$ | Larvae $\alpha\alpha$ vs. $\beta\beta$ |
| Tyrosine metabolism | Amino acid metabolism | 17 | 5 | 3.1 | 0.035 | 0.173 | Male $\alpha\alpha$ vs. $\beta\beta$ | |
| Amino sugar and nucleotide sugar metabolism | Carbohydrate metabolism | 38 | 4 | 6.0 | 0.011 | 0.193 | Adult $\alpha\alpha$ vs. $\beta\beta$ | |
| Glyoxylate and dicarboxylate metabolism | Carbohydrate metabolism | 19 | 7 | 5.1 | 0.014 | 0.146 | Female $\alpha\alpha$ vs. $\beta\beta$ | Adult $\alpha\alpha$ vs. $\beta\beta$ |
| Galactose metabolism | Carbohydrate metabolism | 25 | 6 | 3.1 | 0.039 | 0.180 | Male $\alpha\alpha$ vs. $\beta\beta$ | |
| Starch and sucrose metabolism | Carbohydrate metabolism | 27 | 8 | 3.7 | 0.008 | 0.083 | Male $\alpha\alpha$ vs. $\beta\beta$ | |
| Oxidative phosphorylation | Energy metabolism | 32 | 5 | 4.1 | 0.004 | 0.057 | Male $\alpha\alpha$ vs. $\beta\beta$ | |
| Fanconi anemia pathway | Genetic information processing (replication and repair) | 13 | 7 | 4.0 | 0.010 | 0.150 | Adult $\alpha\alpha$ vs. $\beta\beta$ | |
| Sphingolipid metabolism | Lipid metabolism | 25 | 3 | 5.8 | 0.007 | 0.146 | Female $\alpha\alpha$ vs. $\beta\beta$ | |
| Ether lipid metabolism | Lipid metabolism | 18 | 4 | 3.2 | 0.028 | 0.172 | Male $\alpha\alpha$ vs. $\beta\beta$ | |
| Fatty acid degradation | Lipid metabolism | 28 | 18 | 6.8 | 0.001 | 0.060 | Male $\alpha\alpha$ vs. $\beta\beta$ | Adult $\alpha\alpha$ vs. $\beta\beta$ |
| Fatty acid elongation | Lipid metabolism | 14 | 4 | 7.0 | 0.001 | 0.060 | Male $\alpha\alpha$ vs. $\beta\beta$ | Adult $\alpha\alpha$ vs. $\beta\beta$ |
| Glycerophospholipid metabolism | Lipid metabolism | 49 | 6 | 4.1 | 0.004 | 0.057 | Male $\alpha\alpha$ vs. $\beta\beta$ | |
| One carbon pool by folate | Metabolism of co-factors and vitamins | 11 | 6 | 5.2 | 0.014 | 0.146 | Female $\alpha\alpha$ vs. $\beta\beta$ | |
| Folate biosynthesis | Metabolism of co-factors and vitamins | 29 | 5 | 4.2 | 0.003 | 0.057 | Male $\alpha\alpha$ vs. $\beta\beta$ | Adult $\alpha\alpha$ vs. $\beta\beta$, Female $\alpha\alpha$ vs. $\beta\beta$ |
| Purine metabolism | Nucleotide metabolism | 118 | 42 | 6.5 | 0.000 | 0.000 | Adult $\alpha\alpha$ vs. $\beta\beta$ | Male $\alpha\alpha$ vs. $\beta\beta$ |
| Pyrimidine metabolism | Nucleotide metabolism | 75 | 6 | 3.6 | 0.021 | 0.162 | Adult $\alpha\alpha$ vs. $\beta\beta$ | |
| Longevity regulating pathway - multiple species | Organismal system (aging) | 41 | 3 | 3.5 | 0.023 | 0.162 | Adult $\alpha\alpha$ vs. $\beta\beta$ | Male $\alpha\alpha$ vs. $\beta\beta$ |
| Phototransduction - fly | Organismal system (Sensory system) | 26 | 6 | 4.0 | 0.005 | 0.057 | Male $\alpha\alpha$ vs. $\beta\beta$ | |
| FoxO signaling pathway | Signal transduction | 46 | 6 | 3.3 | 0.031 | 0.176 | Adult $\alpha\alpha$ vs. $\beta\beta$ | |
| Neuroactive ligand-receptor interaction | Signalling molecules | 9 | 7 | 5.3 | 0.011 | 0.146 | Female $\alpha\alpha$ vs. $\beta\beta$ | |

**Table 3** - Functional pathways exhibiting subnetworks of genes interacting with each other and differentially expressed between karyotypes or sexes.
| Pathway | Category | Network Size | Subnetwork Size | Subnetwork Score | P-value | Q-value | Analysis |
| --- | --- | --- | --- | --- | --- | --- | --- |
| Alanine, aspartate and glutamate metabolism | Amino acid metabolism | 26 | 7 | 4.4 | 0.011 | 0.059 | Sex |
| Arginine and proline metabolism | Amino acid metabolism | 28 | 4 | 6.8 | 0.000 | 0.000 | Sex |
| Drug metabolism - cytochrome P450 | Xenobiotics biodegradation and metabolism | 20 | 8 | 3.7 | 0.028 | 0.101 | Sex |
| Drug metabolism - other enzymes | Xenobiotics biodegradation and metabolism | 33 | 5 | 3.6 | 0.033 | 0.110 | Sex |
| Folate biosynthesis | Metabolism of co-factors and vitamins | 29 | 4 | 3.9 | 0.021 | 0.084 | Sex |
| Galactose metabolism | Carbohydrate metabolism | 25 | 7 | 3.8 | 0.022 | 0.084 | Sex |
| Glutathione metabolism | Amino acid metabolism | 37 | 8 | 6.2 | 0.000 | 0.000 | Sex |
| Glycerophospholipid metabolism | Lipid metabolism | 49 | 7 | 3.4 | 0.046 | 0.140 | Sex |
| Glycolysis / Gluconeogenesis | Carbohydrate metabolism | 36 | 4 | 6.9 | 0.000 | 0.000 | Sex |
| Insect hormone biosynthesis | Metabolism of terpenoids and polyketides | 24 | 8 | 3.5 | 0.038 | 0.113 | Sex |
| Longevity regulating pathway - multiple species | Organismal system (aging) | 41 | 8 | 3.6 | 0.035 | 0.111 | Sex |
| Phototransduction - fly | Organismal system (Sensory system) | 26 | 7 | 4.5 | 0.009 | 0.056 | Sex |
| Purine metabolism | Nucleotide metabolism | 118 | 9 | 8.6 | 0.000 | 0.000 | Sex |
| Pyruvate metabolism | Carbohydrate metabolism | 27 | 4 | 6.9 | 0.000 | 0.000 | Sex |
| Taurine and hypotaurine metabolism | Metabolism of other amino acids | 9 | 3 | 4.0 | 0.017 | 0.083 | Sex |
| Valine, leucine and isoleucine degradation | Amino acid metabolism | 32 | 3 | 3.9 | 0.019 | 0.084 | Sex |

Taken together, these GO terms and the gene networks analysis reveal a clear and strong association with development and metabolism/energy storage; and cell cycle metabolism and genetic information processing, respectively. Overall, more terms for the effect of karyotype were associated with the male data set compared to the female data set (GO: 2 terms vs. 1 term, Signet: 15 pathways vs. 8 pathways) although this is not surprising given the difference in the number of differentially expressed genes. These associations between inversion karyotype and metabolism and development are corroborated by the large phenotypic effects of *Cf-Inv(1)*, which results in strong size and developmental time differences in males but not females [31, 35].

There were fewer terms associated with the larvae. Overall, the signal in larvae was very weak and we only identified one pathway significantly differing between genotypes: thiamine metabolism, which is associated with digestion. This is not surprising as larvae stop feeding before pupation [52] and αα males develop 1.2-15x more slowly than ββ males. It should be noted that our larval samples were almost certainly in different stages of development as we standardized by time rather than stage. Work in *Drosophila melanogaster* shows that thiamine is critical for pupation [53] further underlining that the differences we observe are likely partially linked to differences in developmental stage.

### Combining genomic and transcriptomic studies facilitates the identification of candidate genes

By combining our gene expression results with results from a previous study that identified environmentally associated SNP outliers [27], we were also able to identify a small group of strong candidate genes for local adaptation. We compared the position of 997 transcripts that were differentially expressed between karyotypes in one of our 6 contrasts (adult αα vs. ββ, adult male αα vs. ββ, adult female αα vs. ββ, larvae αα vs. ββ, larvae αβ vs. ββ, larvae αα vs. αβ) with 1,526 outlier SNPs identified as being associated with biotic and abiotic characteristics of the wrackbed, as these factors have been found to be significant selective forces on *Cf-Inv(1)*[29, 35, 39]. We found 86 differentially-expressed transcripts that mapped within 5 kb of an environmentally associated SNP. Randomly subsampling our tested transcripts 10,000 times indicated that the expected overlap should only be 42 ± 0.06 transcripts. This is likely due to the linkage disequilibrium created by the inversion, running this test using only transcripts that mapped to *Cf-Inv(1)* generated an expectation closer to the observed value (expected overlap: 67 ± 0.06, actual: 70). Of our 86 overlapping transcripts, 55 were associated with one of two principal components that described seaweed composition of the wrackbed habitat while 44 were associated with abiotic characteristics of the wrackbed such as depth, temperature and salinity. There was some overlap, 13 transcripts were associated with both wrackbed composition and climate. All of the transcripts associated with abiotic characteristics were located in *Cf-Inv(1)*. In contrast, 15/55 transcripts associated with seaweed composition were located in other places in the genome. Full information on these loci can be found in Supplemental Tables 4 and 5.

The wrackbed composition represents a major selective force both on *Cf-Inv(1)* as well as on *C. frigida* as a whole. Flies raised on *Laminaria* spp. are larger and in better condition than flies raised on *Fucus spp.* although this effect is strongest in αα and αβ males [30]. These effects are likely tied directly to the microbial community of these algae, which forms the base of the *C. frigida* larval diet; *Fucus spp.* supports large numbers of *Flavobacterium* whereas *Pseudomonas spp.* are more common on *Laminaria spp.* [54, 55]. Thus, we expect some candidate genes to be related to either digestion or growth. Within our 55 candidates we found several loci relating to digestive processes, such as carbonic anhydrase 5A which helps regulate pH of the midgut in *Drosophila melanogaster* [56] and trypsin, a crucial digestive enzyme [57]. As with the signet analysis, we also uncovered genes relating to the cessation of larval feeding and the onset of pupation, suggesting that the timing of this transition is a major factor underlying the size difference between αα and ββ males rather than differences in larval growth rate. In insects, two of the major modulators of feeding behavior are neuropeptide F (*npf*) and serotonin (5-HT) [58] [59]. In older non-feeding *Drosophila* larvae, *npf* is downregulated [60] and one potential mediator of this is tetrahydrobiopterin (BH4), a fat derived metabolite that suppresses the release of *npf* from *npf* neurons [61]. Among our candidates was pterin-4-alpha-carbinolamine dehydratase (*Pcd*), which is involved in the recycling of BH4 and thus increasing levels of BH4. In our data, *Pcd* was upregulated in ββ larvae and ββ males: it could suppress *npf* and thus feeding behavior leading to earlier pupation. 5-HT is another a major regulator of feeding behavior and increased levels of 5-HT in the gut of *Drosophila melanogaster* enhance larval feeding behavior [59]. Among our candidates was 5-hydroxytryptamine receptor 1 (*HT1R*) which was upregulated in αα males, potentially increasing feeding behavior.

Abiotic characteristics are harder to associate with gene function than seaweed composition but we did find an abundance of genes involved in pupation, cuticle hardening, and eclosion such as *LGR5* and *LCR15* [62], eclosion hormone [63], and *ChT* [64]. Development time in *C. frigida* is highly plastic and is affected by temperature and density as well as karyotype [38]. As wrackbeds are ephemeral habitats there is likely strong selection on these traits as well. Overall, these results provide some initial insights and putative candidates for further exploration. Furthermore, it is clear that many of the traits are likely polygenic and highly complex. While merging transcriptomic and genomic datasets provides an excellent first step to narrow down candidates, more work, especially functional validation, needs to be done to differentiate between adaptive and linked variation.

## Conclusions

Abundant evidence indicates that chromosomal inversions are key genomic factors in eco-evolutionary processes because of their multifarious impacts on genome structure, recombination and regulation [8, 10]. However, few studies have made progress towards dissecting the mechanistic pathways that enable inversions to shape evolutionary trajectories. Using a transcriptomic approach in the seaweed fly *Coelopa frigida* revealed that the impact of *Cf-Inv(1)* was conditional and differed between males, females, and larvae. Males showed a stronger effect of *Cf-Inv(1)* than females. Overall, most of the differentially expressed genes were *cis*-regulated for karyotype, but not for sex effects. Interestingly, genes where the effect of *Cf-Inv(1)* was more constant were more likely to be *cis*-regulated than genes whose differential expression was conditional. These results suggest that *trans* regulation may be important for conditional gene expression in inversions. Combining our results with genomic data uncovered candidate variants in the inversion that may underlie mechanistic pathways that determine critical phenotypes in particular the cessation of larval feeding. Overall, our results highlight the complex effects of inversion polymorphisms on gene expression across contexts and the benefit of combining transcriptomic and genomic approaches in the study of inversions.

## METHODS

### Rearing and crosses

Larvae of *C. frigida* for breeding were collected from the field in April/May 2017 from Skeie, Norway (58.69733, 5.54083), Østhassel, Norway (58.07068, 6.64346), Ystad, Sweden (55.425, 13.77254), and Smygehuk, Sweden (55.33715, 13.35963). Larvae were also collected from Skadbergsanden, Norway (58.45675, 5.91407) in June 2016. See Figure 1 for all sampling locations. All larvae were brought back live to the Tjärnö Marine Laboratory in Strömstad, Sweden where they were raised to adulthood at 25°C.

We generated an αα line from Skeie and a ββ line from each population (see supplemental methods for details). Six days after the creation of these lines two replicates of 3 larvae each from each line were flash frozen in liquid nitrogen and stored at −80°C until extraction. Larvae were always stored as groups of 3 henceforth referred to as larval pools. The adults that emerged from these lines were used to make subsequent crosses within and between karyotypes and populations to generate αβ and ββ larvae (see Supplemental Table 5 for the crossing scheme). Adults were then flash frozen individually in liquid nitrogen and stored at −80°C until extraction. All experimental crosses were set up in a 50 mL tube with a sponge for aeration and 4 g *Saccharina latissima* and 2 g *Fucus spp*. Six days after the creation of these crosses one larval pool from each cross was flash frozen in liquid nitrogen and stored at −80°C until extraction. All larval pools and adults were processed at the same time of day (+/− 1 hour) to reduce variation. We were able to get larval pools from two successful crosses per cross type. We also generated a ontogeny series to ensure a comprehensive transcriptome (Supplemental Note).

### RNA extraction, Library preparation and sequencing

RNA from all samples was extracted following a TriZOL protocol (Supplementary Note). Only flies from our lab lines and crosses were sequenced: 2 larval pools per line (1 αα and 4 ββ lines) and 2 larval pools from each subsequent cross type (see Supplemental Table 5 for the crossing scheme). We also sequenced 3 Skeie αα adult males, 3 Skeie αα adult females, 5 Skeie ββ adult males, 2 Skeie ββ adult females, 3 Skadbergsanden ββ adult females, and 1 Ystad ββ adult female. We chose these samples to bias towards parents of the larval samples and endeavored to get a good distribution of genotypes. However, we were severely limited by RNA quality. All of these samples were submitted to SciLifeLab in Uppsala, Sweden for library preparation and sequencing. RNA was purified with Agencourt RNA clean XP before library preparation. Library preparation was done with the TruSeq stranded mRNA library preparation kit including polyA selection. Samples were sequenced on a NovaSeq S1 flowcell with 100 bp paired end reads (v1 sequencing chemistry).

### Transcriptome assembly

We only used samples from the geographically close populations Skeie and Østhassel to construct our transcriptome to limit genetic variation between samples. Individual assemblies for 2 of the Skeie αα adult males, 2 of the Skeie αα adult females, 2 of the Skeie ββ adult males, 2 of the Skeie ββ adult females, both of the Østhassel ontogenetic pools spanning 0-348 hours of development (as a single assembly), both of the Skeie αα larval pools (as a single assembly), and both of the Skeie ββ larval pools (as a single assembly) were done using Trinity v2.9.1 (11 assemblies in total)[65]. Prior to assembly, all reads were trimmed and adaptors removed using cutadapt 2.3 with Python 3.7.2 [66]. All assemblies were run through TransRate 1.0.1 [41], a quality assessment tool for *de novo* transcriptomes that looks for artifacts, such as chimeras and incomplete assembly, and provides individual transcript and overall assembly scores. We retained all transcripts from each assembly classified by TransRate as ‘good’. These contigs were then merged using CD-hit 4.8.1 [67] with a sequence identity threshold of 0.95, a word size of 10, and local sequence alignment coverage for the longer sequence at 0.005. Finally, the transcriptome was mapped to the genome assembly [27] using GMAP 2018-07-04 [68]. The mapping coordinates for each transcript were extracted and in the event that two transcripts mapped to the same coordinates, only the longer transcript was retained. The mapping coordinates of all transcripts were retained for use in further analyses. The final transcriptome was annotated using the Trinotate pipeline with the Uniprot/Swiss-Prot and Pfam databases (Downloaded on June 25^th^, 2020) [69].

### Differential expression analysis

We used DESeq2 1.26.0 to determine which transcripts were differentially expressed between karyotypes and sexes [70]. The reads from all samples were trimmed and the adaptors were removed using cutadapt 2.3 with Python 3.7.2 [66]. The trimmed reads were then aligned to the reference transcriptome using bowtie2 2.3.5.1 [71] and quantified using RSEM [72]. The resulting genes.results files were prepared for use in DESeq2 using the Trinity script abundance_estimates_to_matrix.pl [65]. These files were used as input for DESeq2 1.26.0 implemented in R [70]. Adults and larvae were analysed separately and normalization was done by DESeq2. We removed all transcripts where the total count of reads (across all individuals) was less than 10. We also removed a single sample (Skeie ββ larvae pool 1) as hierarchical clustering using a distance matrix revealed that this sample was an extreme outlier. In DESeq2 our model for adults included both karyotype and sex and their interaction, while the model for larvae included karyotype and population. We did not include population in the adult model as 13/17 samples came from the Skeie population. We further split adult males and females and analyzed them separately. Conventional thresholds (log2 fold change > 2, adjusted p-value (FDR) < 5%) were used to identify differentially expressed transcripts. We tested for gene ontology enrichment in our different sets of results using topGO [73]with the elim algorithm and the Fisher exact test implemented in R [70]. Manhattan distance matrices for all subgroups (males, females, and larvae) were calculated using the dist() function in R and PERMANOVA results were calculated using Adonis2 in the vegan package [45]. Note that karyotype was always used at the first term as terms are added sequentially and models differed between subgroups.

### Gene sub-network analysis

To investigate the effect of inversion on expression in genes involved in common biological pathways, we performed a gene network analysis designed to detect polygenic selection using the R package *signet* [51]. This method defines sub-networks of genes that interact with each other, because they are known to be involved in the same biological pathway in the KEGG database, and present similar patterns attributed to selection; for example covariation in expression levels. For this analysis we used the *Drosophila melanogaster* KEGG database and thus focused on the transcripts that matched a gene in Flybase (13,586 out of 26,239). Variation of expression levels between genotypes were analysed in a multivariate framework with redundancy analysis (RDA), with and without sex as covariate, and scaled to a z-score such that individual transcript scores have a mean of 0 and a standard deviation of 1 (following [74]). Following the recommendations of the *signet* procedure, each pathway of the KEGG database was parsed to score gene sub-networks using 10,000 iterations of simulated annealing. A null distribution of sub-network scores was generated by random sampling to create 10,000 sub-networks of variable sizes. We consider as significant pathways with a higher score than the null distribution, that is, with a p-value below 0.05, and a false-discovery-rate (q-value) of 0.20.

### Overlap with genomic results

We combined our data with previously published population genomic data to identify loci that may contribute to local adaptation. Briefly, in our previous work, 16 populations of *C. frigida* were sampled along latitudinal and ecological gradients and sequenced at the whole-genome level, and the association between SNPs and environmental variation was tested using a combination of two genotype-environment association methods (LFMM2 and Baypass) [27]. Using our mapping coordinates we identified transcripts located <5kb from an outlier SNP defined by both of these association methods and differentially expressed between genotypes in at least one of our analyses.

### Allele specific expression

We used our set of αβ larvae to search for transcripts that showed allele specific expression (ASE). RNA from each of our samples was mapped to our reference genome using bowtie2 2.3.5.1 [71]. The alignment files were sorted and read groups were added using Picard 2.10.3 (http://broadinstitute.github.io/picard/). The resulting files were indexed with samtools [75] and SNPs were called using bcftools [75]. We took the conservative approach of only examining loci that were fixed different between arrangements. SNPs were filtered by mean depth (>5), maximum percentage of missing samples (25%), and F_ST_ between α and β = 1, using vcftools [76]. We further retained only SNPs that had observations from at least 3 individuals. To test for allele specific expression we used the ASEP package [46] implemented in R [70]. This package utilizes multi-individual information and accounts for multi-SNP correlations within the transcripts. Using ASEP we performed a one-condition analysis to detect gene-level ASE and corrected for multiple testing using the Benjamini and Hochberg (1995) method implemented in R with ‘p.adjust.’ [77]. We considered contigs with an adjusted P-value < 0.1 to be significant.

## Acknowledgements

We thank I. Fragata, L. Bernatchez and C. Rougeux for support and advice on analysis and figures. E.L.B. was supported by a Marie Skłodowska-Curie fellowship 704920 – ADAPTIVE INVERSIONS and gratefully acknowledges funding from Helge Ax:son Johnsons Stiftelse. C.M. was supported by fellowship from Fonds de Recherche du Québec (FRQS260724, FRQNT200125) and a Banting postdoctoral fellowship (#162647). H.P. gratefully acknowledges funding from The Swedish Foundation for Strategic Environmental Research MISTRA (grant no. 2013/75). M.W. gratefully acknowledges funding from the Swedish Research Council (grant no. 2012-03996_VR).

## Author Contributions

E.L.B, R.K.B, K.J. and M.W conceived the idea. E.L.B carried out the breeding experiments and labwork. E.L.B and C.M analyzed the data. E.L.B and H.P provided financing. E.L.B and M.W. wrote the manuscript with input from all authors.

## REFERENCES

1. Twyford AD, Friedman J. Adaptive divergence in the monkey flower *Mimulus guttatus* is maintained by a chromosomal inversion. Evolution. 2015;69(6):1476–86.

2. Westram AM, Faria R, Johannesson K, Butlin R. Using replicate hybrid zones to understand the genomic basis of adaptive divergence. Molecular Ecology. 2021.

3. Kirkpatrick M, Barton N. Chromosome inversions, local adaptation and speciation. Genetics. 2006;173(1):419–34.

4. Knief U, Forstmeier W, Pei YF, Ihle M, Wang DP, Martin K, et al. A sex-chromosome inversion causes strong overdominance for sperm traits that affect siring success. Nat Ecol Evol. 2017;1(8):1177–84. doi: 10.1038/s41559-017-0236-1. PubMed PMID: WOS:000417188600025.

5. Lemaitre C, Braga MDV, Gautier C, Sagot MF, Tannier E, Marais GAB. Footprints of Inversions at Present and Past Pseudoautosomal Boundaries in Human Sex Chromosomes. Genome Biology and Evolution. 2009;1:56–66. doi: 10.1093/gbe/evp006. PubMed PMID: WOS:000275269200007.

6. Peichel CL, Ross JA, Matson CK, Dickson M, Grimwood J, Schmutz J, et al. The master sex-determination locus in threespine sticklebacks is on a nascent Y chromosome. Curr Biol. 2004;14(16):1416–24. doi: DOI 10.1016/j.cub.2004.08.030. PubMed PMID: WOS:000223586900019.

7. Ayala D, Guerrero RF, Kirkpatrick M. Reproductive isolation and local adaptation quantified for a chromosome inversion in a malaria mosquito. Evolution. 2013;67(4):946–58.

8. Hoffmann AA, Rieseberg LH. Revisiting the impact of inversions in evolution: From population genetic markers to drivers of adaptive shifts and speciation? Annual review of ecology, evolution, and systematics. 2008;39:21–42.

9. Lowry DB, Willis JH. A widespread chromosomal inversion polymorphism contributes to a major life-history transition, local adaptation, and reproductive isolation. Plos Biol. 2010;8(9). doi: 10.1371/journal.pbio.1000500. PubMed PMID: WOS:000282279200019.

10. Wellenreuther M, Bernatchez L. Eco-evolutionary genomics of chromosomal inversions. Trends Ecol Evol. 2018;33(6):427–40. doi: 10.1016/j.tree.2018.04.002. PubMed PMID: WOS:000432462300007.

11. Küpper C, Stocks M, Risse JE, Dos Remedios N, Farrell LL, McRae SB, et al. A supergene determines highly divergent male reproductive morphs in the ruff. Nature genetics. 2016;48(1):79–83.

12. Lamichhaney S, Fan G, Widemo F, Gunnarsson U, Thalmann DS, Hoeppner MP, et al. Structural genomic changes underlie alternative reproductive strategies in the ruff (Philomachus pugnax). Nature genetics. 2016;48(1):84–8.

13. Joron M, Frezal L, Jones RT, Chamberlain NL, Lee SF, Haag CR, et al. Chromosomal rearrangements maintain a polymorphic supergene controlling butterfly mimicry. Nature. 2011;477(7363):203–6.

14. Noor MA, Grams KL, Bertucci LA, Reiland J. Chromosomal inversions and the reproductive isolation of species. Proceedings of the National Academy of Sciences. 2001;98(21):12084–8.

15. Berdan EL, Blanckaert A, Butlin RK, Bank C. Deleterious mutation accumulation and the long-term fate of chromosomal inversions. Plos Genet. 2021;17(3):e1009411. doi: https://doi.org/10.1371/journal.pgen.1009411.

16. Crow T, Ta J, Nojoomi S, Aguilar-Rangel MR, Rodríguez JVT, Gates D, et al. Gene regulatory effects of a large chromosomal inversion in highland maize. Plos Genet. 2020;16(12):e1009213.

17. Kozak GM, Brennan RS, Berdan EL, Fuller RC, Whitehead A. Functional and population genomic divergence within and between two species of killifish adapted to different osmotic niches. Evolution. 2014;68(1):63–80.

18. Renaut S, Nolte AW, Rogers SM, Derome N, Bernatchez L. SNP signatures of selection on standing genetic variation and their association with adaptive phenotypes along gradients of ecological speciation in lake whitefish species pairs (Coregonus spp.). Molecular ecology. 2011;20(3):545–59.

19. Pardo - Diaz C, Salazar C, Jiggins CD. Towards the identification of the loci of adaptive evolution. Methods in Ecology and Evolution. 2015;6(4):445–64.

20. Shanta O, Noor A, Sebat J. The effects of common structural variants on 3D chromatin structure. BMC genomics. 2020;21(1):1–10.

21. Lupiáñez DG, Kraft K, Heinrich V, Krawitz P, Brancati F, Klopocki E, et al. Disruptions of topological chromatin domains cause pathogenic rewiring of gene-enhancer interactions. Cell. 2015;161(5):1012–25.

22. Lavington E, Kern AD. The effect of common inversion polymorphisms In (2L) t and In (3R) Mo on patterns of transcriptional variation in Drosophila melanogaster. G3: Genes, Genomes, Genetics. 2017;7(11):3659–68.

23. Lettice LA, Daniels S, Sweeney E, Venkataraman S, Devenney PS, Gautier P, et al. Enhancer - adoption as a mechanism of human developmental disease. Human mutation. 2011;32(12):1492–9.

24. Huang W, Carbone MA, Magwire MM, Peiffer JA, Lyman RF, Stone EA, et al. Genetic basis of transcriptome diversity in Drosophila melanogaster. Proceedings of the National Academy of Sciences. 2015;112(44):E6010–E9.

25. Said I, Byrne A, Serrano V, Cardeno C, Vollmers C, Corbett-Detig R. Linked genetic variation and not genome structure causes widespread differential expression associated with chromosomal inversions. Proceedings of the National Academy of Sciences. 2018;115(21):5492–7.

26. Fuller ZL, Haynes GD, Richards S, Schaeffer SW. Genomics of natural populations: how differentially expressed genes shape the evolution of chromosomal inversions in Drosophila pseudoobscura. Genetics. 2016;204(1):287–301.

27. Merot C, Berdan E, Cayuela H, Djambazian H, Ferchaud A-L, Laporte M, et al. Chromosomal rearrangements represent modular cassettes for local adaptation across different geographic scales. bioRxiv. 2020.

28. Aziz JB. Investigations into chromosomes 1, 2 and 3 of *Coelopa frigida* (Fab.): Thesis; 1975.

29. Day TH, Dawe C, Dobson T, Hillier PC. A chromosomal inversion polymorphism in Scandinavian populations of the seaweed fly, *Coelopa frigida*. Hereditas. 1983;99(1):135–45. doi: 10.1111/j.1601-5223.1983.tb00738.x.

30. Edward DA. Habitat composition, sexual conflict and life history evolution in *Coelopa frigida*: University of Sterling; 2008.

31. Butlin RK, Read IL, Day TH. The effects of a chromosomal inversion on adult size and male mating success in the seaweed fly, *Coelopa frigida*. Heredity. 1982;49(1):51–62.

32. Gilburn AS, Day TH. Sexual dimorphism, sexual selection and the α β chromosomal inversion polymorphism in the seaweed fly, *Coelopa frigida*. Proceedings: Biological Sciences. 1994;257(1350):303–9.

33. Mérot C, Llaurens V, Normandeau E, Bernatchez L, Wellenreuther M. Balancing selection via life-history trade-offs maintains an inversion polymorphism in a seaweed fly. Nature communications. 2020;11(1):1–11.

34. Butlin R, Day T. Adult size, longevity and fecundity in the seaweed fly, Coelopa frigida. Heredity. 1985;54(1):107–10.

35. Mérot C, Berdan EL, Babin C, Normandeau E, Wellenreuther M, Bernatchez L. Intercontinental karyotype – environment parallelism supports a role for a chromosomal inversion in local adaptation in a seaweed fly. P Roy Soc B-Biol Sci. 2018;285(1881). doi: ARTN 20180519 10.1098/rspb.2018.0519. PubMed PMID: WOS:000436565200008.

36. Berdan EL, Rosenquist H, Larson K, Wellenreuther M. Inversion frequencies and phenotypic effects are modulated by the environment: insights from a reciprocal transplant study in *Coelopa frigida*. Evol Ecol. 2018;32(6):683–98. doi: 10.1007/s10682-018-9960-5. PubMed PMID: WOS:000450510900007.

37. Butlin RK. The maintenance of an inversion polymorphism in *Coelopa frigida*: University of Nottingham; 1983.

38. Butlin RK, Day TH. The effect of larval competition on development time and adult size in the seaweed fly, *Coelopa frigida*. Oecologia. 1984;63(1):122–7.

39. Butlin RK, Day TH. Environmental correlates of inversion frequencies in natural populations of seaweed flies (*Coelopa frigida*). Heredity. 1989;62(2):223–32.

40. Wellenreuther M, Rosenquist H, Jaksons P, Larson W. Local adaptation along an environmental cline in a species with an inversion polymorphism. Journal of Evolutionary Biology. 2017;30(6):1068–77.

41. Smith-Unna R, Boursnell C, Patro R, Hibberd JM, Kelly S. TransRate: reference-free quality assessment of de novo transcriptome assemblies. Genome research. 2016;26(8):1134–44.

42. Simão FA, Waterhouse RM, Ioannidis P, Kriventseva EV, Zdobnov EM. BUSCO: assessing genome assembly and annotation completeness with single-copy orthologs. Bioinformatics. 2015;31(19):3210–2.

43. Dunn D, Crean C, Wilson C, Gilburn A. Male choice, willingness to mate and body size in seaweed flies (Diptera: Coelopidae). Animal Behaviour. 1999;57(4):847–53.

44. Crean C, Gilburn A. Sexual selection as a side-effect of sexual conflict in the seaweed fly, Coelopa ursina (Diptera: Coelopidae). Animal Behaviour. 1998;56(6):1405–10.

45. Dixon P. VEGAN, a package of R functions for community ecology. Journal of Vegetation Science. 2003;14(6):927–30.

46. Fan J, Hu J, Xue C, Zhang H, Susztak K, Reilly MP, et al. ASEP: Gene-based detection of allele-specific expression across individuals in a population by RNA sequencing. Plos Genet. 2020;16(5):e1008786.

47. Fraser HB. Gene expression drives local adaptation in humans. Genome research. 2013;23(7):1089–96.

48. Knight JC. Allele-specific gene expression uncovered. Trends in Genetics. 2004;20(3):113–6.

49. Snoek BL, Sterken MG, Bevers RP, Volkers RJ, van’t Hof A, Brenchley R, et al. Contribution of trans regulatory eQTL to cryptic genetic variation in C. elegans. BMC genomics. 2017;18(1):1–15.

50. Signor SA, Nuzhdin SV. The evolution of gene expression in cis and trans. Trends in Genetics. 2018;34(7):532–44.

51. Gouy A, Daub JT, Excoffier L. Detecting gene subnetworks under selection in biological pathways. Nucleic Acids Research. 2017;45(16):e149–e.

52. Chown SL, Gaston KJ. Body size variation in insects: a macroecological perspective. Biol Rev. 2010;85(1):139–69.

53. Sannino DR, Dobson AJ, Edwards K, Angert ER, Buchon N. The Drosophila melanogaster gut microbiota provisions thiamine to its host. MBio. 2018;9(2).

54. Bolinches J, Lemos ML, Barja JL. Population dynamics of heterotrophic bacterial communities associated withFucus vesiculosus andUlva rigida in an estuary. Microbial ecology. 1988;15(3):345–57.

55. Laycock R. The detrital food chain based on seaweeds. I. Bacteria associated with the surface of Laminaria fronds. Marine Biology. 1974;25(3):223–31.

56. Overend G, Luo Y, Henderson L, Douglas AE, Davies SA, Dow JA. Molecular mechanism and functional significance of acid generation in the Drosophila midgut. Scientific reports. 2016;6(1):1–11.

57. Wu D-D, Wang G-D, Irwin DM, Zhang Y-P. A profound role for the expansion of trypsin-like serine protease family in the evolution of hematophagy in mosquito. Molecular biology and evolution. 2009;26(10):2333–41.

58. Fadda M, Hasakiogullari I, Temmerman L, Beets I, Zels S, Schoofs L. Regulation of feeding and metabolism by neuropeptide F and short neuropeptide F in invertebrates. Frontiers in endocrinology. 2019;10:64.

59. Neckameyer WS. A Trophic Role for Serotonin in the Development of a Simple Feeding Circuit. Developmental Neuroscience. 2010;32(3):217–37. doi: 10.1159/000304888.

60. Wu Q, Wen T, Lee G, Park JH, Cai HN, Shen P. Developmental control of foraging and social behavior by the Drosophila neuropeptide Y-like system. Neuron. 2003;39(1):147–61.

61. Kim D-H, Shin M, Jung S-H, Kim Y-J, Jones WD. A fat-derived metabolite regulates a peptidergic feeding circuit in Drosophila. Plos Biol. 2017;15(3):e2000532.

62. Mendive FM, Van Loy T, Claeysen S, Poels J, Williamson M, Hauser F, et al. Drosophila molting neurohormone bursicon is a heterodimer and the natural agonist of the orphan receptor DLGR2. FEBS letters. 2005;579(10):2171–6.

63. Krüger E, Mena W, Lahr EC, Johnson EC, Ewer J. Genetic analysis of Eclosion hormone action during Drosophila larval ecdysis. Development. 2015;142(24):4279–87.

64. Hamid R, Hajirnis N, Kushwaha S, Saleem S, Kumar V, Mishra RK. Drosophila Choline transporter non-canonically regulates pupal eclosion and NMJ integrity through a neuronal subset of mushroom body. Developmental biology. 2019;446(1):80–93.

65. Haas BJ, Papanicolaou A, Yassour M, Grabherr M, Blood PD, Bowden J, et al. De novo transcript sequence reconstruction from RNA-seq using the Trinity platform for reference generation and analysis. Nature protocols. 2013;8(8):1494–512.

66. Martin M. Cutadapt removes adapter sequences from high-throughput sequencing reads. 2011. 2011;17(1):3. Epub 2011-08-02. doi: 10.14806/ej.17.1.200.

67. Fu L, Niu B, Zhu Z, Wu S, Li W. CD-HIT: accelerated for clustering the next-generation sequencing data. Bioinformatics. 2012;28(23):3150–2.

68. Wu TD, Watanabe CK. GMAP: a genomic mapping and alignment program for mRNA and EST sequences. Bioinformatics. 2005;21(9):1859–75.

69. Grabherr MG, Haas BJ, Yassour M, Levin JZ, Thompson DA, Amit I, et al. Full-length transcriptome assembly from RNA-Seq data without a reference genome. Nature biotechnology. 2011;29(7):644–52.

70. Love MI, Huber W, Anders S. Moderated estimation of fold change and dispersion for RNA-seq data with DESeq2. Genome biology. 2014;15(12):550.

71. Langmead B, Salzberg SL. Fast gapped-read alignment with Bowtie 2. Nature methods. 2012;9(4):357.

72. Li B, Dewey CN. RSEM: accurate transcript quantification from RNA-Seq data with or without a reference genome. BMC bioinformatics. 2011;12(1):323.

73. Alexa A, Rahnenfuhrer J. topGO: enrichment analysis for gene ontology. R package version. 2010;2(0):2010.

74. Rougeux C, Gagnaire PA, Praebel K, Seehausen O, Bernatchez L. Polygenic selection drives the evolution of convergent transcriptomic landscapes across continents within a Nearctic sister species complex. Molecular ecology. 2019;28(19):4388–403.

75. Li H. A statistical framework for SNP calling, mutation discovery, association mapping and population genetical parameter estimation from sequencing data. Bioinformatics. 2011;27(21):2987–93.

76. Danecek P, Auton A, Abecasis G, Albers CA, Banks E, DePristo MA, et al. The variant call format and VCFtools. Bioinformatics. 2011;27(15):2156–8.

77. Benjamini Y, Hochberg Y. Controlling the false discovery rate: a practical and powerful approach to multiple testing. Journal of the Royal statistical society: series B (Methodological). 1995;57(1):289–300.

