## Supplemental Figures for "A large chromosomal inversion shapes gene expression in seaweed flies (*Coelopa frigida*)"

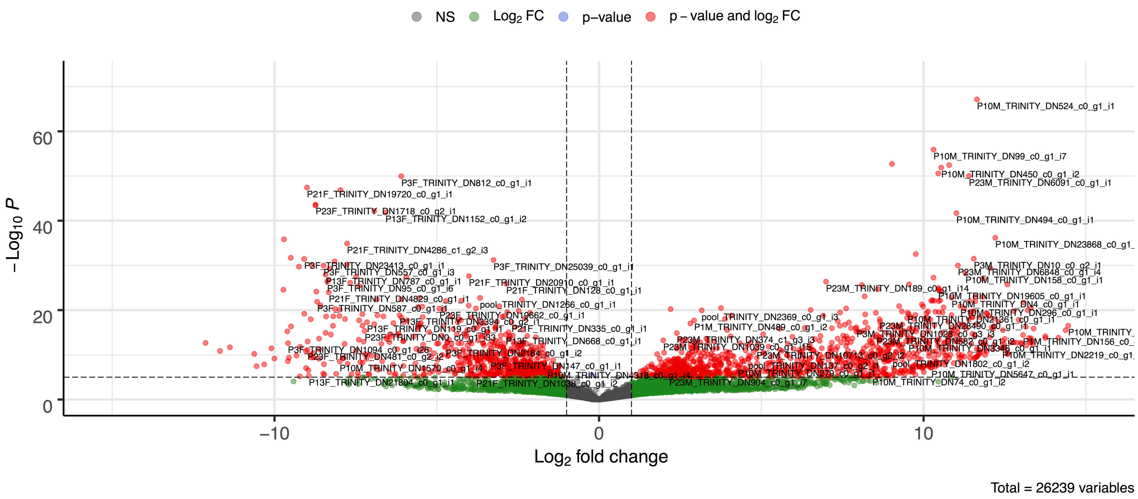


Supplemental Figure 1 - Differential expression by sex in adults. Positive values indicate higher expression in males and negative values indicate higher expression in females. Each dot represents a single gene and is colored to indicate significance. Grey - not significant, Green - significant by $log_2$ fold change (>2), Blue - significant by p-value (<0.001), Red - significant by $log_2$ fold change and p-value. Figure made with EnhancedVolcano implemented in R [1].

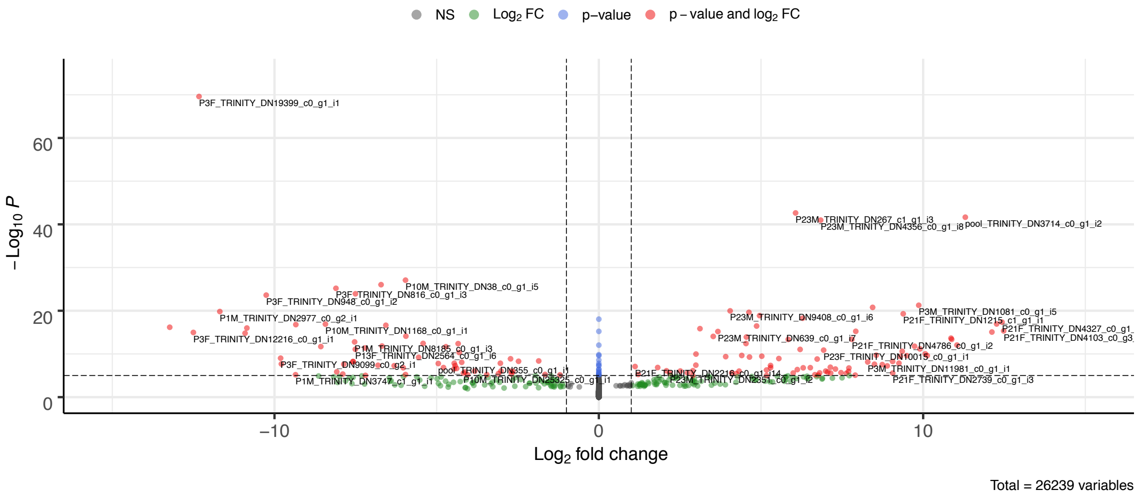


Supplemental Figure 2 - Differential expression by karyotype in adults. Positive values indicate higher expression in ββ and negative values indicate higher expression in αα. Each dot represents a single gene and is colored to indicate significance. Grey - not significant, Green - significant by $log_2$ fold change (>2), Blue - significant by p-value (<0.001), Red - significant by $log_2$ fold change and p-value. Figure made with EnhancedVolcano implemented in R [1].

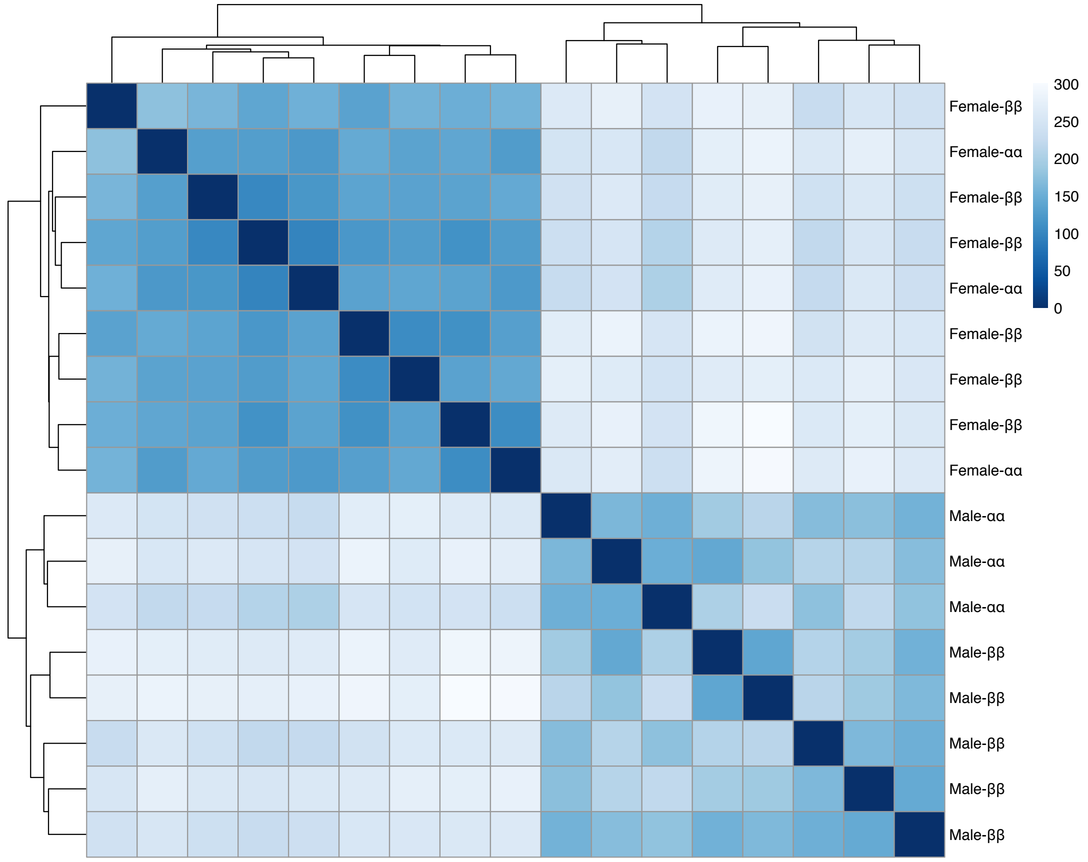


<u>Supplemental Figure 3</u> - Heatmap of Euclidian sample distances in adults. Samples are labeled with sex and karyotype. Color indicates distance, with darker blues being more similar. Figure made with pheatmap implemented in R [2].

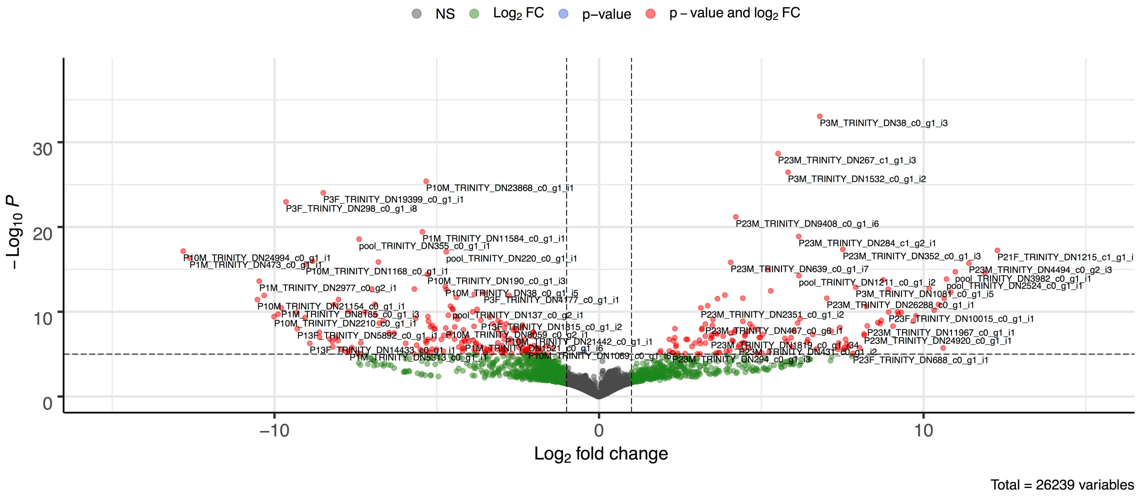


Supplemental Figure 4 - Differential expression by karyotype in adult males. Positive values indicate higher expression in ββ and negative values indicate higher expression in αα. Each dot represents a single gene and is colored to indicate significance. Grey - not significant, Green - significant by $log_2$ fold change (>2), Blue - significant by p-value (<0.001), Red - significant by $log_2$ fold change and p-value. Figure made with EnhancedVolcano implemented in R [1].

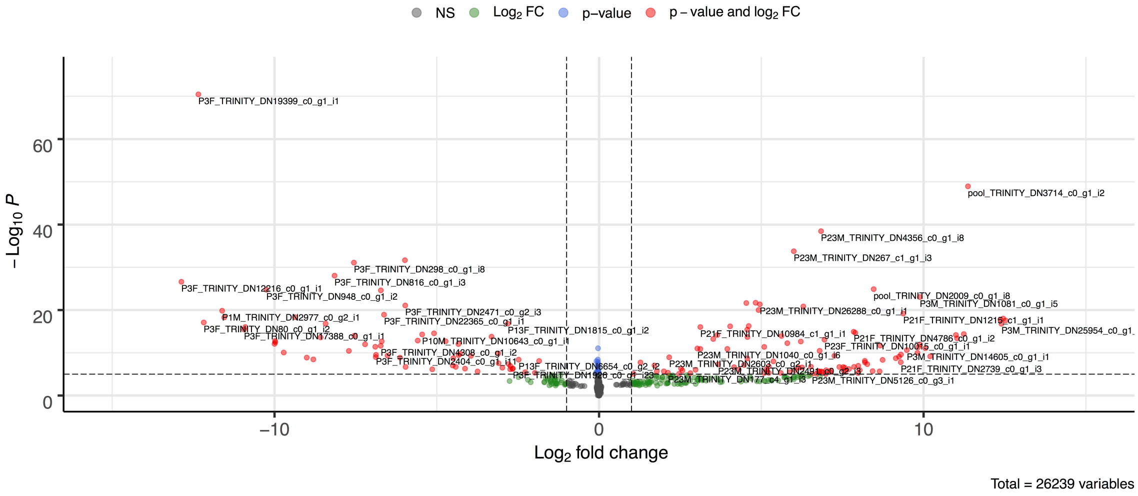


Supplemental Figure 5 - Differential expression by karyotype in adult females. Positive values indicate higher expression in ββ and negative values indicate higher expression in αα. Each dot represents a single gene and is colored to indicate significance. Grey - not significant, Green - significant by $log_2$ fold change (>2), Blue - significant by p-value (<0.001), Red - significant by $log_2$ fold change and p-value. Figure made with EnhancedVolcano implemented in R [1].

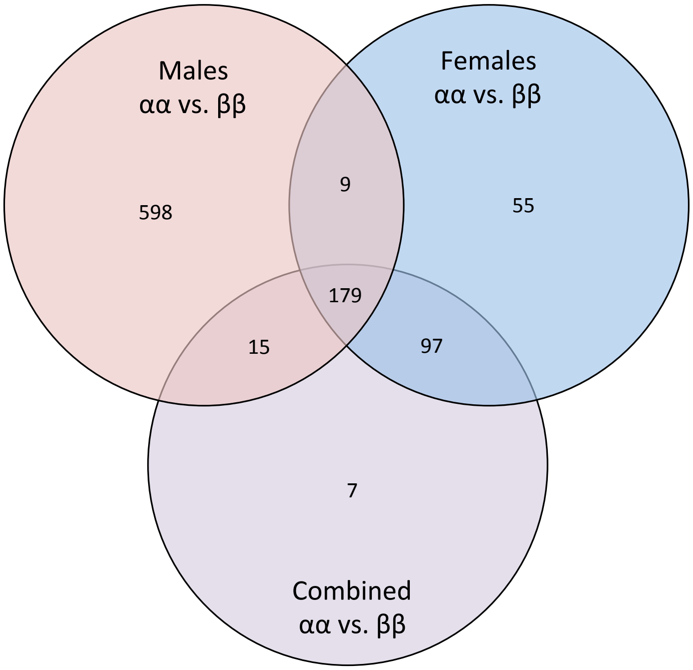


Supplemental Figure 6 - Overlap between differential expression test sets. Pictured is the number of significantly differentially expressed transcripts that overlapped between our 3 test sets: male αα vs. ββ, female αα vs. ββ, combined (i.e. both sexes) αα vs. ββ.

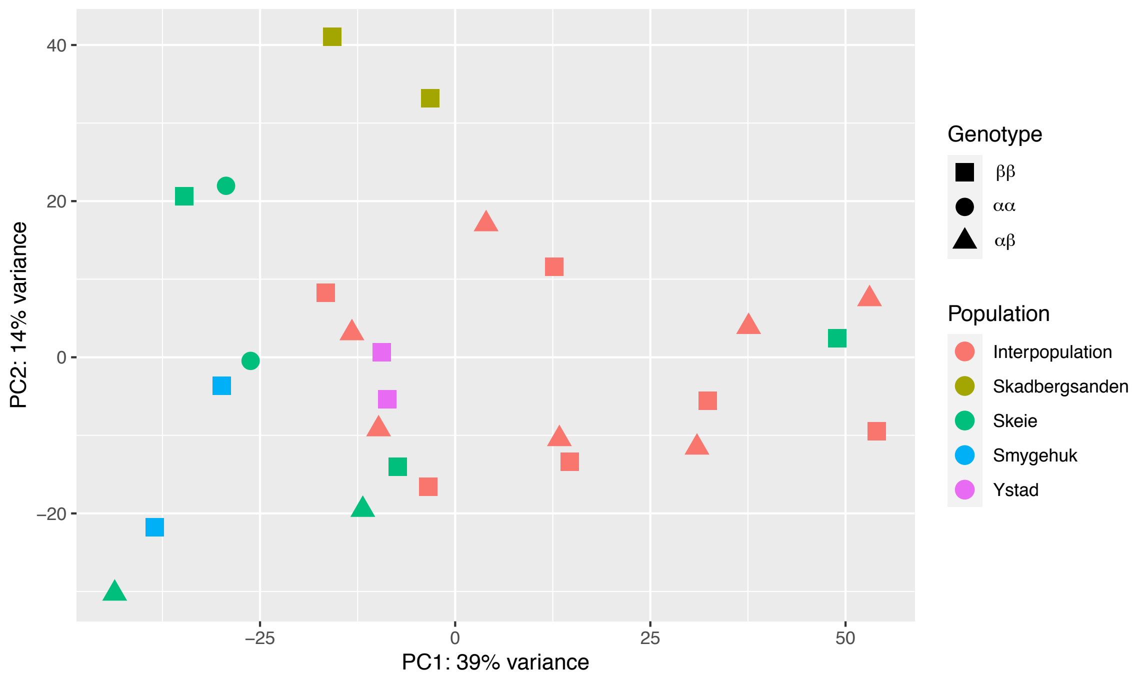


Supplemental Figure 7 - Expression variation by population in larvae. Points are colored by population (Interpopulation - red, Skadbergsanden - mustard, Skeie - green, Smygehuk - blue, and Ystad-purple) and shaped according to karyotype (αα-triangle, αβ-square, ββ-circle). Interpopulation refers to larvae that were generated by interpopulation crosses, see supplemental table 1 for cross types.

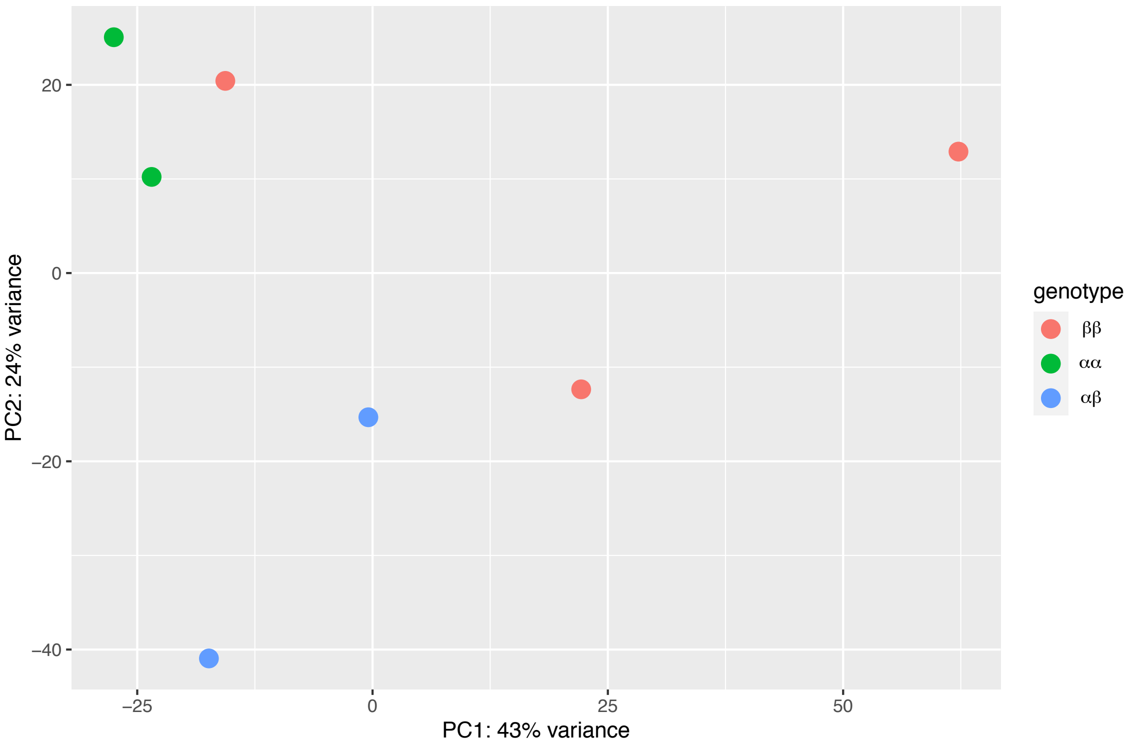


Supplemental Figure 8 - Principal component analysis (PCA) of expression variation in larvae from Skeie. Points are colored by karyotype (αα- green, αβ-blue, ββ-red).

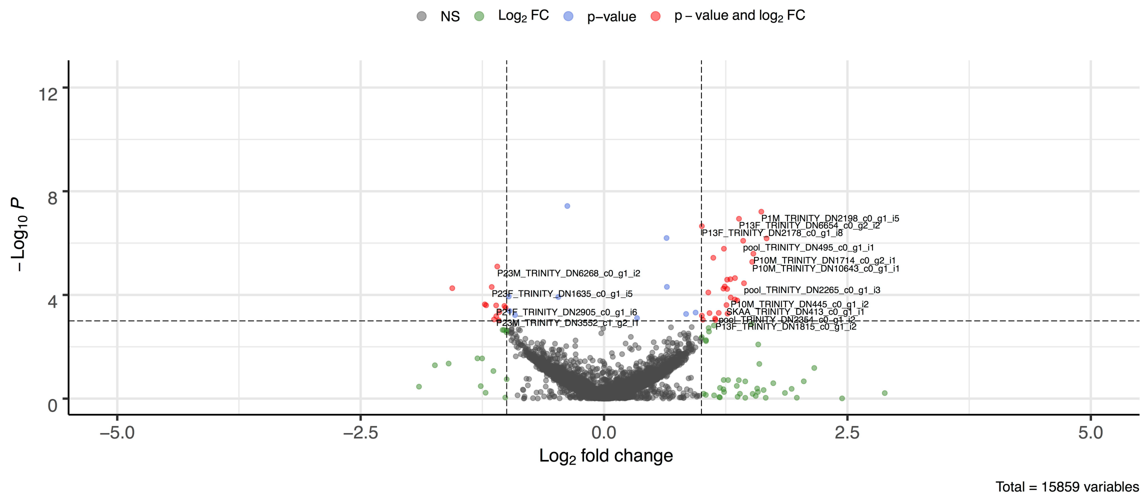


Supplemental Figure 9 - Differential expression by genotype in larvae. Positive values indicate higher expression in αβ and negative values indicate higher expression in ββ. Each dot represents a single gene and is colored to indicate significance. Grey - not significant, Green - significant by $\log_2$ fold change (>2), Blue - significant by p-value (<0.001), Red - significant by $\log_2$ fold change and p-value. Figure made with EnhancedVolcano implemented in R [1].

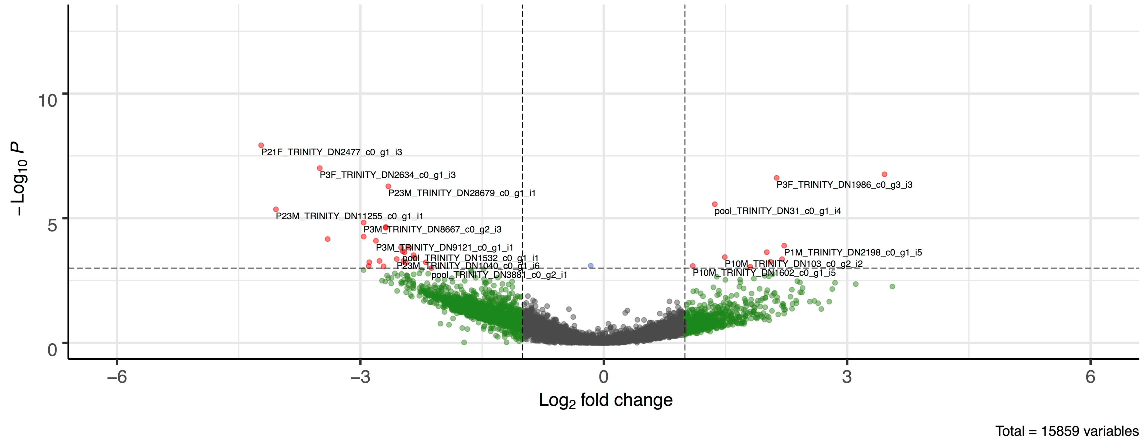


Supplemental Figure 10 - Differential expression by genotype in larvae. Positive values indicate higher expression in αα and negative values indicate higher expression in ββ. Each dot represents a single gene and is colored to indicate significance. Grey - not significant, Green - significant by $\log_2$ fold change (>2), Blue - significant by p-value (<0.001), Red - significant by $\log_2$ fold change and p-value. Figure made with EnhancedVolcano implemented in R [1].

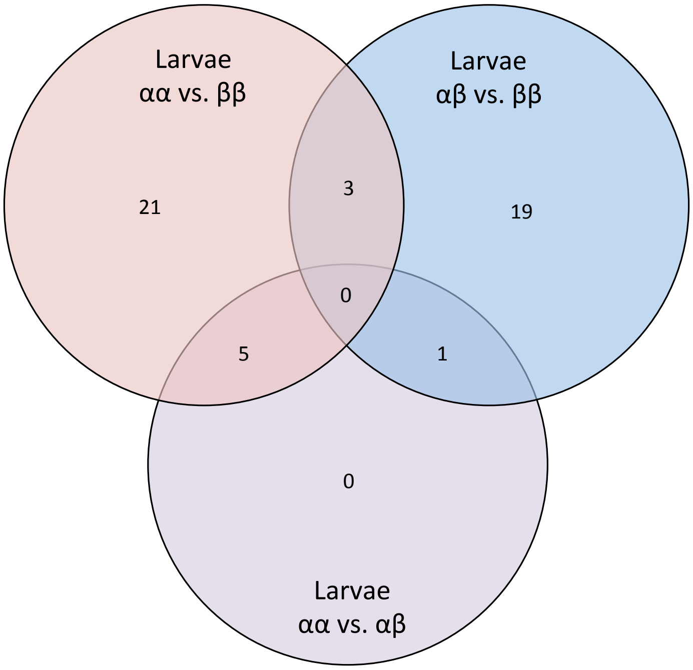


Supplemental Figure 11 - Overlap between differential larval expression test sets. Pictured is the number of significantly differentially expressed transcripts that overlapped between our 3 test sets: αα vs. ββ, αβ vs. ββ, and αα vs. αβ.

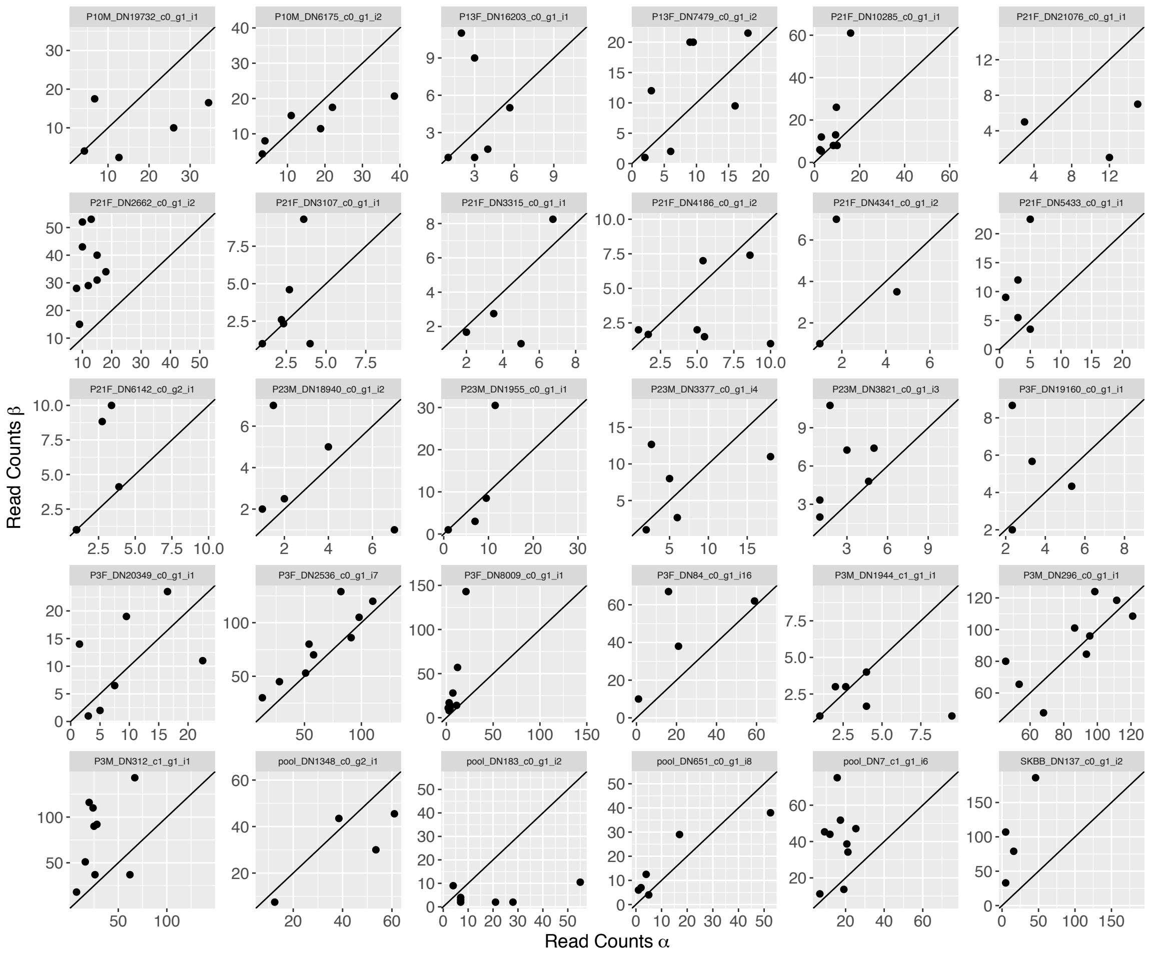


<u>Supplemental Figure 12</u> - Transcripts with significant allele specific expression. Each plot is for a single transcript where each dot represents a single individual averaged over all SNPs in that transcript. A 1:1 line is provided for context.

1. Blighe K, Rana S, Lewis M. EnhancedVolcano: Publication-ready volcano plots with enhanced colouring and labeling. R package version. 2019;1(0).
2. Kolde R, Kolde MR. Package 'pheatmap'. 2015.
