## Supplemental Text for "A large chromosomal inversion shapes gene expression in seaweed flies (*Coelopa frigida*)"

### METHODS

#### Rearing and crosses

All lines were maintained in the lab at 25°C with a 12:12 light:dark cycle and fed on a 1:1 mix of *Saccharina latissima* and *Fucus spp* We generated both αα and ββ lines for the Skeie population but only ββ lines from all other populations. For all crosses described below we used virgin adults. Virgin adults were obtained by putting individual pupae into small aerated Eppendorf tubes with a small amount of cotton soaked in 0.5% mannitol. Twenty four to forty eight hours after eclosion adults were placed in a refrigerator (4-5°C) and kept there until used. To create specific karyotype lines, one of the hind legs of each adult was removed for genotyping. DNA extraction and genotyping for a SNP in complete linkage disequilibrium with *Cf-Inv(1)* was performed as described in [32]. From Skeie, 3 separate αα x αα and 3 separate ββ x ββ crosses were obtained, from all other populations 3 separate ββ x ββ crosses were obtained. These crosses were used to generate our lines. For each line, thirty individuals (15 ♀ and 15 ♂), combined over the 3 founder crosses, were introduced to new 0.8L aerated containers containing 90 g *Saccharina latissima* and 45 g *Fucus spp.*

#### Time series

To ensure that we generated a comprehensive transcriptome, adults from Østhassel were used to generate a development time series. Sixty males and sixty females (20 individuals/pot/sex) were introduced to three replicate pots with 50% *Saccharina latissima* and 50% *Fucus spp.* substrate. They were left for 24 hours to lay eggs after which they were removed. At this time eggs were sampled and the pots were sampled every subsequent 48 hours for 17 days until adults had been observed for 3 subsequent time points. For each sample, 3-6 individuals were pooled depending on availability. If multiple stages were present at a single time point (e.g., larvae and pupae) then 3-6 individuals per stage were pooled in the sample. Samples were stored in RNAlater in -20°C until extraction. We extracted RNA from 6 time points (0 hours, 48 hours, 96 hours, 192 hours, 288 hours, and 384 hours). For each time point we separately extracted samples from two of the three replicate pots. Concentration of these extractions was measured using a Qubit and equal amounts of RNA from each time point from each of the two samples were pooled to make two parallel pools. Each pool covered the same time and stage distribution, but created from different samples.

#### RNA extraction

RNA from all samples was extracted following a TriZOL protocol. Briefly 500 µl of TriZOL was added to each sample and then the sample was homogenized in a shaker using glass beads. The sample was then incubated at room temperature for 5 minutes after which it was centrifuged at 12,000 rcf 4°C for 10 minutes. The supernatant was transferred to a new tube and 100 µl of chloroform was added. After shaking, the tube was incubated at room temperature for 3 minutes after which it was centrifuged at 10,000 rcf at 4°C for 15 minutes. The upper aqueous phase was transferred to a new tube and 100 µl of phenol and 100 µl of chloroform were added. After shaking the sample was centrifuged for 7 minutes at 10,000 rcf at 4°C. Then, 500 µl of isopropanol was added and the sample was incubated for 10 minutes at room temperature. After a centrifugation of 10 minutes at 12,000 rcf at 4°C the pellet was washed with 500 µl of 75% EtOH, centrifuged at 7,500 rcf at 4°C for 5 minutes, and air-dried for 10 minutes. Samples were re-suspended in 50 µl of nuclease-free $H_20$. After at least 24 hours, all samples were further cleaned using the Zymo RNA Clean & Concentrator Kit following manufacturer's instructions.

#### RESULTS

##### Single individual transcriptomes

The 11 single sample assemblies we generated had between 6,583 and 48,558 transcripts (Supplemental Table 2). Assemblies made from larvae had on average fewer and shorter transcripts (mean # - 7,939, mean N50 - 938) than assemblies made from adults (mean # - 36,881, mean N50 - 2,179). We input a total of 300,113 transcripts into CD-HIT and found 38,142 clusters. The representative sequence from each cluster was mapped to version 1.0 of the genome and we removed 2,143 transcripts that mapped to the same coordinates as a longer transcript. We retained all transcripts that did not map to the genome at all (1,233).
